# Performance of three freely available methods for extracting white matter hyperintensities: FreeSurfer, UBO Detector and BIANCA

**DOI:** 10.1101/2020.10.17.343574

**Authors:** Isabel Hotz, Pascal Frédéric Deschwanden, Franziskus Liem, Susan Mérillat, Spyridon Kollias, Lutz Jäncke

**Affiliations:** Division of Neuropsychology, Department of Psychology, University of Zurich, Switzerland; University Research Priority Program (URPP), Dynamics of Healthy Aging, University of Zurich, Zurich, Switzerland; Department of Neuroradiology, University Hospital Zurich, Zurich, Switzerland

**Keywords:** White matter hyperintensities, Automated segmentation, MRI, Healthy aging, Validation

## Abstract

White matter hyperintensities (WMH) of presumed vascular origin are frequently found in MRIs of healthy older adults. WMH are also associated with aging and cognitive decline. Here, we compared and validated three freely available algorithms for WMH extraction: FreeSurfer, UBO Detector, and BIANCA (Brain Intensity AbNormality Classification Algorithm) using a longitudinal dataset comprising MRI data of cognitively healthy older adults (baseline *N* = 231, age range 64–87 years). As reference we manually segmented WMH in T1w, 3D FLAIR, 2D FLAIR images. These manual segmentations were then used to assess the segmentation accuracy of the different automated algorithms. Further, we assessed the relationships of WMH volume estimates provided by the algorithms with Fazekas scores and age. FreeSurfer underestimated the WMH volumes and scored worst in Dice Similarity Coefficient (DSC = 0.434) but its WMH volumes strongly correlated with the Fazekas scores (*r_s_* = 0.73). BIANCA accomplished the highest DCS (0.602) with 3D FLAIR images. However, the relations with the Fazekas scores were only moderate, especially in the 2D FLAIR images (*r_s_* = 0.41), and many outlier WMH volumes were detected when exploring within-person trajectories (2D FLAIR: ~30%). UBO Detector performed similarly to BIANCA in DSC with both modalities and reached the best DSC in 2D FLAIR (0.531) without requiring a tailored training dataset. In addition, it achieved very high associations with the Fazekas scores (2D FLAIR: *r_s_* = 0.80).

In summary, our results emphasize the importance of carefully contemplating the choice of the WMH segmentation algorithm and MR-modality.

## 1. Introduction

According to the STandards for Reporting Vascular changes on nEuroimaging (STRIVE), white matter hyperintensities (WMH) of presumed vascular origin appear as hyperintense signal abnormalities on T2-weighted magnetic resonance images (MRI), can appear as isointense or hypointense on T1-weighted sequences, and vary in diameter (Wardlaw et al., 2013). The fluid-attenuated inversion recovery (FLAIR) sequence is generally the most sensitive structural MRI sequence for detecting WMH (Wardlaw et al., 2013). They are a common finding in MR images of older adults (Wardlaw, Valdés Hernández, & Muñoz-Maniega, 2015) and considered as a marker of cerebrovascular disease (CVD), which, in turn, is a major cause of morbidity and mortality in older age and triggers cognitive decline (Baker et al., 2012). Even in healthy older adults WMH are associated with reduced cognitive, perceptual and motor abilities (Di Stadio et al., 2020; Gunning-Dixon & Raz, 2000; Pinter et al., 2017). More generally, the presence of WMH increases the risk of global functional loss (Inzitari et al., 2009), stroke, dementia, and death (Debette & Markus, 2010). Therefore, precise and reliable WMH quantification using the most sensitive MR sequences is of key importance.

In the context of clinical diagnostics, the Fazekas scale (Fazekas, Chawluk, Alavi, Hurtig, & Zimmerman, 1987), the Scheltens scale (Scheltens et al., 1993), and the age-related white matter changes scale (ARWMC) (Wahlund et al., 2001) are commonly used to visually assess the severity and progression of WMH. However, despite the relatively simply use of visual rating scales they unfortunately do not provide true quantitative data, are time-consuming to obtain (Mäntylä et al., 1997), are not sensitive enough to assess longitudinal changes in WMH (D. M. J. van den Heuvel et al., 2006) because of significant ceiling and floor effects (Mäntylä et al., 1997). Compared to such scales, volumetric measurements are more reliable and more sensitive to detect age-related changes in longitudinal studies of WMH (T. L. A. van den Heuvel et al., 2016). Especially for investigations of cognitively healthy samples, where the expected WMH volume increases over time is rather small, we therefore need fast automated methods that provide the estimated volume rather than a score (Frey et al., 2019; Prins & Scheltens, 2015).

Segmenting the WMH manually is extremely time-consuming and can become prohibitively expensive, and is therefore not feasible when considering the current trend towards big data (i.e., datasets with a large *N* and multiple time points of data acquisition). Hence, accurate automatic methods for WMH lesion volume quantification are highly desirable. To date, there are many different methods for WMH quantification which can be roughly divided into: *Supervised*, when the underlying algorithm is trained using a manually segmented «gold standard» as a reference (Ghafoorian et al., 2017; Van Nguyen, Zhou, & Vemulapalli, 2015), and *unsupervised*, when the method does not rely on a gold standard (Bowles et al., 2016; Cardoso, Sudre, Modat, & Ourselin, 2015; Ye, Zikic, Glocker, Criminisi, & Konukoglu, 2013). Depending on the amount of human intervention required, the methods can further be divided into *automated vs. semi-automated* (Caligiuri et al., 2015). A recent study of Guerrero et al. (2018) provides a comprehensive review of existing methods. All of these methods face the challenge of false-positive and false-negative segmentations as well as different WM lesion loads (usually lower in MS lesions than in WMH of presumed vascular origin) and different WM lesion contrasts (MS lesions are typically brighter and more sharply bordered compared to WMH of presumed vascular origin (Caligiuri et al., 2015; Griffanti et al., 2016)). Co-occurring pathologies (e. g., extensive atrophy) further challenge the methods (Heinen et al., 2019).

Caligiuri and colleagues (2015) compared different existing algorithms including supervised/ unsupervised and automated/semi-automated methods. They found that many of these methods are not freely available, are study and/or protocol specific, and have been validated primarily with small samples. Importantly, there is still no consensus on which algorithm(s) is (are) of good quality and should be applied to detect WMH (Dadar et al., 2017; Frey et al., 2019). Consequently, the methodology of pertinent studies is very heterogeneous and compromises the comparability of such studies. Therefore, the primary goal of our current work is to assess the performance of three freely available WMH extraction methods: FreeSurfer (Fischl, 2012), UBO Detector (Jiang et al., 2018), and BIANCA (Brain Intensity AbNormality Classification Algorithm) (Griffanti et al., 2016).

The FreeSurfer Image Analysis Suite (Fischl, 2012) is a fully automated software of tools for analysis of brain structures using information of T1w images. Although FreeSurfer was not developed specifically for WMH segmentation, it may be a useful option, especially if no FLAIR images were collected in a study. Although single metrics have been provided for the WMH segmentation algorithm of FreeSurfer (Ajilore et al., 2014; Olsson et al., 2013; Samaille et al., 2012; Smith et al., 2011) common accuracy metrics are still lacking. Further, existing validations rely on images with a low WMH load and the algorithm has not yet been validated longitudinally.

UBO Detector is a cluster-based, fully automated pipeline that works without a training dataset and, like BIANCA, relies on the *k*-NN algorithm for quantifying WMH (Jiang et al., 2018). It was validated by the developers themselves, using two datasets – one cross-sectional and the other longitudinal – both with 2D FLAIR images. The groups included older participants with medical conditions such as stroke, transient ischemic attack (TIA), etc. They showed that UBO Detector is a reliable tool for WMH extraction and found strong WMH agreement compared to their manual reference. To date, no further validation study of UBO Detector has been published, but it was used for extracting WMH in three additional studies (d’Arbeloff et al., 2019; Du & Xu, 2019; Taylor et al., 2019).

BIANCA (Brain Intensity AbNormality ClassificationAlgorithm) is a semi-automated, multimodal, supervised method for WMH detection, based on the *k*-nearest neighbor (*k*-NN) algorithm (Griffanti et al., 2016). Besides a global thresholding, BIANCA also offers the LOCally Adaptive Threshold Estimation (LOCATE), a supervised method for identifying optimal local thresholds to apply to the estimated lesion probability map (Sundaresan et al., 2018). BIANCA was validated and optimized cross-sectionally with a «predominantly neurodegenerative» cohort, and with a «predominantly vascular» cohort (Griffanti et al., 2016), and was shown to perform better when compared against the two freely available methods «CASCADE» (Damangir et al., 2012) and «Lesion Segmentation Tool» (P. Schmidt et al., 2012). Ling and colleagues (2018) validated and optimized BIANCA cross-sectionally based on a cohort of patients with CADASIL (Cerebral autosomal dominant arteriopathy with subcortical infarcts and leukoencephalopathy) and concluded that BIANCA is a reliable method to extract extensive WMH loads in these patients. Sundaresan et al. (2018) validated BIANCA by including LOCATE (see above) cross-sectionally on different cohorts with a wide range of WMH loads, and showed that LOCATE contributed to a better segmentation performance, especially in CADASIL patients. So far, BIANCA with global thresholding and BIANCA with the local threshold method LOCATE have not been validated with a longitudinal dataset. LOCATE has not yet been validated using data with low WMH load with a manually segmented reference. For an overview of articles validating FreeSurfer, UBO Detector, and BIANCA, see **Supplementary Table 1.**

In this study, we aim to provide complementary information on the performance of the three automated WMH extraction algorithms depending on different MRI input modalities (T1w, 2D FLAIR + T1w, 3D FLAIR + T1w). We also evaluated differences between the fully-automated FreeSurfer and UBO Detector methods and the semi-automated BIANCA algorithm. To estimate the accuracy of the automated WMH segmentations, we used fully manually segmented WMH as a proxy for true WMH. We refer to these manually segmented WMH as gold standards.

To our knowledge, this is the first study that evaluates the WMH segmentation performance of different methods and the influence of MR image modality using comprehensive longitudinal MRI data of cognitively healthy older adults collected in a single-center study.

With our study we want to answer the following explicit questions:

1. ***Assessment of segmentation accuracy:*** Which algorithm and MR image modality provides estimates that are most consistent with the respective gold standard in terms of:
  1a) established accuracy metrics *and*
  1b) WMH volumes.
2. ***Validations using the whole dataset:*** How do the WMH volume estimates provided by the different algorithms relate to
  2a) the frequently used Fazekas score *and*
  2b) chronological age?

## 2. Methods

### 2.1. Subjects

Longitudinal MRI data were taken from the Longitudinal Healthy Aging Brain Database Project (LHAB; Switzerland) – an ongoing project conducted at the University of Zurich (Zöllig et al., 2011). We used data from the first four measurement occasions (baseline, 1-year follow-up, 2-year follow-up, 4-year follow-up). The baseline LHAB dataset includes data from 232 participants (mean age at baseline: *M* = 70.8, range = 64–87; F:M = 114:118). At each measurement occasion, participants completed an extensive battery of neuropsychological and psychometric cognitive tests and underwent brain imaging. Inclusion criteria for study participation at baseline were age ≥ 64 years, righthandedness, fluent German language proficiency, a score of ≥ 26 points on the Mini Mental State Examination (Folstein, Folstein, & McHugh, 1975), no self-reported neurological disease of the central nervous system and no contraindications to MRI. The study was approved by the ethical committee of the canton of Zurich. Participation was voluntary and all participants gave written informed consent in accordance with the declaration of Helsinki.

For the present analysis, we used participants with structural MRI data with a sample size at baseline of *N* = 231 (mean age at baseline: *M* = 70.8, range = 64–87; F:M = 113:118). At 4-year follow-up, 71.9% of the baseline sample still comprised structural data (*N* = 166, mean age: M = 74.2, range = 68–87; F:M = 76:90). In accordance with previous studies in this field, we ensured that none of the included scans showed intracranial hemorrhages, intracranial space occupying lesions, multiple sclerosis (MS) lesions, large chronic, subacute or acute infarcts, and extreme visually apparent movement artefacts.

### 2.2. MRI data acquisition

MRI data were acquired at the University Hospital of Zurich on a Philips Ingenia 3T scanner (Philips Medical Systems, Best, The Netherlands) using the dsHead 15-channel head coil. T1w and 2D FLAIR structural images were part of the standard MRI battery of the LHAB project, and are therefore available for the most time points (see **Table 1**). T1w images were recorded with a 3D T1w turbo field echo (TFE) sequence, repetition time (TR): 8.18 ms, echo time (TE): 3.799 ms, flip angle (FA): 8°, 160 × 240 × 240 mm^3^ field of view (FOV), 160 sagittal slices, in-plain resolution: 256 × 256, voxel size: 1.0 × 0.94 × 0.94 mm^3^, scan time: ~7:30 min. If two T1w images were available per time point, the images were averaged for further use. The 2D FLAIR image parameters were: TR: 11000 ms, TE: 125 ms, inversion time (TI): 2800 ms, 180 × 240 × 159 mm^3^ FOV, 32 transverse slices, in-plain resolution: 560 × 560, voxel size: 0.43 × 0.43 × 5.00 mm^3^, interslice gap: 1 mm, scan time: ~5:08 min. 3D FLAIR images were recorded for a subsample only. The 3D FLAIR image parameters were: TR: 4800 ms, TE: 281 ms, TI: 1650 ms, 250 × 250 mm FOV, 256 transverse slices, in-plain resolution: 326 × 256, voxel size: 0.56 × 0.98 × 0.98 mm^3^, scan time: ~4:33 min.

**Table 1.** Number of scans (N) broken down for modality (3D T1w, 2D FLAIR, 3D FLAIR) and time point (Tp).

| Modality | Time points |  |  |  | Total N |
| --- | --- | --- | --- | --- | --- |
|  | Tp1 n | Tp2 n | Tp3 n | Tp5 n |  |
| 3D T1-weighted | 231 | 207 | 196 | 166 | 800 |
| 2D FLAIR | 228 | 203 | 174 | 157 | 762 |
| 3D FLAIR | 4 | 46 | 53 | 63 | 166 |

**Table 1** provides an overview of the number of available MRI scans per data acquisition time point (baseline, 1-year follow-up, 2-year follow-up, 4-year follow-up) and image modality (T1w, 2D FLAIR, 3D FLAIR). Please see **Supplementary Figure 1** for an extended overview of the data structure and study design.

### 2.3. Subsets of dataset used for the different algorithms

In this work we used three subsets of the LHAB dataset to validate the algorithms. The subsets differed with respect to the MR image modalities and in the number of scans. UBO Detector and BIANCA extract WMH-related intensity features mainly from the FLAIR images, since FLAIR images provide the best WMH contrast. In addition, T1w images are used for both algorithms. In UBO Detector, T1w images are required (for the segmentation of white matter, gray matter and cerebrospinal fluid tissues). BIANCA allows for additional T1w image input and it has been shown that the additional inclusion of T1w images improved segmentation performance (Griffanti et al., 2016). **Table 2** provides an overview of the subsets used for the different algorithms.

**Table 2.** Name of the dataset subsets (FreeSurfer T1w, UBO 2D, BIANCA 2D, UBO 3D, BIANCA 3D), applied algorithms, input modality/modalities for the different algorithms, and the number (N) of scans per subset.

| Name of the subsets | Algorithm(s) | Input modality/<br>modalities | Total N of scans |
| --- | --- | --- | --- |
| FreeSurfer T1w | FreeSurfer | 3D T1w | 800 |
| UBO 2D<br>BIANCA 2D | UBO Detector & BIANCA | 2D FLAIR + 3D T1w | 762 |
| UBO 3D<br>BIANCA 3D | UBO Detector & BIANCA | 3D FLAIR + 3D T1w | 166 |

### 2.4. Accuracy metrics

We used several metrics for quantifying the segmentation performance of the different algorithms. These metrics provide information about the degree of overlap, the degree of resemblance, and the volumetric agreement when comparing (a) gold standards manually segmented by multiple operators and (b) algorithm outputs with gold standards. The equations of these metrics are listed in **Table 3**.

**Table 3.** The following metrics were used to determine the agreements between the operators (Inter-Operator) and between the outcomes of the algorithms and the gold standards (Validation).

| Metrics | Formulas<br>(Inter-Operator) | Formulas<br>(Validation) |
| --- | --- | --- |
| Hausdorff distance for the 95th percentile (H95) | $d(A, B) = {}^{95}K_{a \in A}^{th} d(a, B) \dagger$ | |
| Dice Similarity Coefficient (DSC) | $\frac{2 * V_A \cap B}{V_A + V_B}$ | $\frac{2 * TP}{FP + 2 * TP + FN}$ |
| Detection Error Rate (DER)<br>(All Clusters without intersection $V_A \cap B$ divided by MTA ‡) | $\frac{V_A + V_B}{MTA}$ | $\frac{2 * (FP + FN)}{FP + FN + 2 * TP}$ |
| Outline Error Rate (OER)<br>(All Clusters with intersection $V_A \cap B$ divided by MTA) | $\frac{V_A + V_B - 2 * V_A \cap B}{MTA}$ | $\frac{2 * (FP + FN)}{FP + FN + 2 * TP}$ |
| Sensitivity = True positive ratio (TPR) | | $\frac{TP}{TP + FN}$ |
| False positive ratio (FPR) | | $\frac{FP}{FP + TN}$ |

Notes: $\dagger {}^{95}K_{a \in A}^{th}$ represents the $K^{th}$ ranked distance such that $K/N_a = 95\%$ (Dubuisson & Jain, 1994). ‡ MTA = mean total area, area of rater A and area of rater B divided by 2 (Wack et al., 2012).

#### Spatial overlap metrics

The Dice Similarity Coefficient (DSC) (Dice, 1945) provides information about the overlap agreement between two segmentations and it is perhaps the most established metric in evaluating the accuracy of WMH segmentation methods. It is defined as two times the union of the selected voxels divided by the sum of the selected voxels by each of the raters or algorithms. However, since the DSC depends on the lesion load (the higher the lesion load, the higher the DSC), it is difficult to evaluate operators or automated segmentation methods against each other if assessed on different sets of scans with different lesion loads (Wack et al., 2012).

Extending the DSC, the Outline Error Rate (OER) and Detection Error Rate (DER) are independent of lesion burden (Wack et al., 2012). In these metrics, the sum of false positive (FP) and false negative (FN) voxels is split, depending on whether an intersection occurred or not. The sum is then divided by the mean total area (MTA) of the two operators to obtain a ratio with the DER as a metric of errors without intersection, and with the OER as a metric of errors with an intersection. OER and DER can also be adopted for the comparison between gold standards and algorithms (see Table 3). We calculated and reported the sensitivity or true positive ratio (TPR). The specificity (also referred to as recall) was not declared as it is equal to 1 – false positive ratio (FPR), which we reported.

#### Spatial distance metric

A different approach to validate an image segmentation is based on distance. The Hausdorff distance, is a shape comparison method and can be used to evaluate how far apart subsets of a metric space are from each other (Beauchemin, Thomson, & Edwards, 1998; Huttenlocher, Klanderman, & Rucklidge, 1993). It represents the maximum distance of a point in one set to the nearest point in the other set (Shonkwiler, 1991). To avoid problems with noisy segmentations we used the modified Hausdorff distance for the 95th percentile (H95) (Huttenlocher et al., 1993).

#### Volumetric agreement

We additionally calculated the Interclass Correlation Coefficient (ICC). The ICC is a measurement that reflects not only the degree of correlation, but also the agreement between two measurements based on mean squares (Koo & Li, 2016). For comparisons with a gold standard, we used the «unit» single (ICC(3,1)) and for the comparison without a gold standard, we used the equation with a pooled average (ICC(3, *k*)) (Koo & Li, 2016; McGraw & Wong, 1996; Shrout & Fleiss, 1979).

### 2.5. Manual WMH segmentation

From the entire longitudinal dataset, which comprises four time points of data acquisition, we selected subjects for whom T1w, 2D FLAIR, and 3D FLAIR scans were available for a given time point. From those, we randomly selected a subsample of 16 subjects for manual segmentation while ensuring that the images of the selected subjects covered a mixed range of WMH loads. To do so, we used individual Fazekas scores. The subsample for manual segmentation had a medium Fazekas score on average. Although Griffanti et al. (2016) recommend a training dataset with only high WMH load, in their study, Ling et al. (2018) achieved better results with a training dataset with a mixed WMH load than with a training dataset with only a high WMH load. Therefore, and because a mixed range of WMH loads represents our dataset more appropriately, we have chosen our sampling strategy (for data structure and study design see **Supplementary Figure 1**).

In total, 48 MR images (16 subjects × 3 image modalities) were manually segmented for WMH, resulting in 48 binary masks with the values *0* for no WMH and *1* for WMH. We refer to these WMH segmentations as gold standards given that they represent strong proxies for the true WMH load.

#### Segmentation of FLAIR images

Three operators (O1, O2, O3) completely manually segmented the WMH on 16 3D FLAIR images and on ten 2D FLAIR images in all three planes (sagittal, coronal, axial) on a MacBook Pro 13-inch with a Retina Display with a screen resolution of 2560 × 1600 pixels, 227 pixels per inch with full brightness intensity to obtain comparable data using FSLeyes (McCarthy, 2018). The segmentations were carried out independently resulting in three different masks per segmented image. Because of the very good preceding segmentation-inter-operator reliabilities, only one operator (O2) segmented the remaining six 2D FLAIR images to achieve the same number (*n* = 16) of gold standards for all modalities. To ensure high segmentation quality, these six WMH masks were peer-reviewed by O1 and O3. Any discrepancies were discussed between all operators and one of the authors (*S.K*.), a neuroradiology professor with over 30 years of experience in diagnosing cerebral MR images. The three manually segmented WMH masks (of the three operators) of the same subject were displayed as overlays in FSLeyes in order to evaluate the mask agreement across these three operators (voxel value *1.0*: all three operators classified the voxel as WMH; voxel value 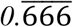 two operators classified the voxel as WMH; 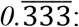 one operator classified the voxel as WMH (see **Figure 1**). Each mask overlay was then revised voxel-by-voxel by consensus in the presence of all operators (O1, O2, O3), and converted back to a binary mask to serve as gold standard. The between-operator disagreements mostly regarded voxels at the WMH borders. The resulting masks were shown to *S.K*. and corrected in case of mistakes.

**Figure 1.**
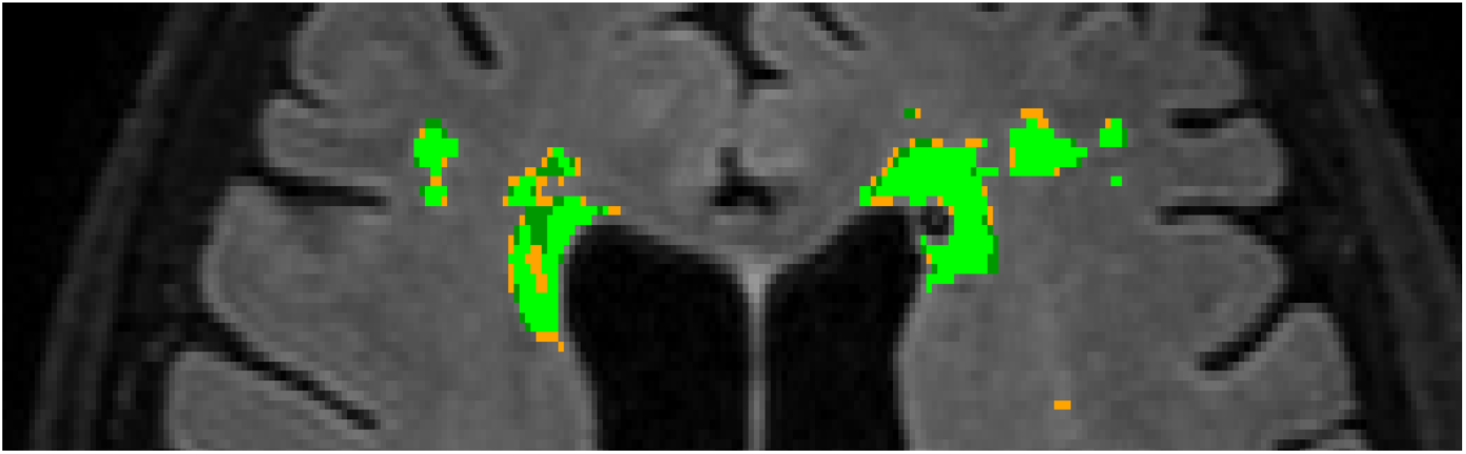
Section of an overlay of three masks – one per operator – named as «mean mask» in a 3D FLAIR image on an axial plane with the different values displayed in different colors per operator (light green: all three operators classified the voxel as WMH (voxel value 1.0); dark green: two operators classified the voxel as WMH (voxel value 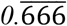); orange: one operator classified the voxel as WMH (voxel value 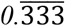).

#### Segmentation of T1w images

The fully manual segmentation of the 16 T1w MR images was apportioned among two operators (O1, O2). Since T1w images provide the lowest WMH contrast, we used the non-segmented FLAIR images from the same subject and time point in case of uncertainties, for example when the contrast for DWMH was very poor or when lacunes penetrated WMH. Nevertheless, we wanted to be influenced by the FLAIR images as little as possible, and therefore consulted the FLAIR images as rarely as possible. In addition, *S.K*. was consulted in case of ambiguities and O1 discussed all images voxel-by-voxel with O3 at the end of the segmentation.

#### Validation of the gold standards

The mean DSC between all three operators for the 3D (*n* = 16) and 2D FLAIR images (*n* = 10 subjects) was 0.73 and 0.67, respectively. A mean DSC of 0.7 (Anbeek, Vincken, van Osch, Bisschops, & van der Grond, 2004; Caligiuri et al., 2015) is considered as a good segmentation. As expected, due to the lower surface-to-volume ratio, the DSC was lower for images with a low WMH load than for images with a high WMH load (Wack et al., 2012). In images with a low WMH load, a DSC above 0.5 is still considered as a very good agreement (Dadar et al., 2017). Our average DSC results in both modalities (3D and 2D FLAIR images) were higher than 0.7 for medium WMH load, and higher than 0.6 for low WMH load, which can be considered as excellent agreement. The reliability of the volumetric agreement between the segmentations of the 3D and 2D FLAIR images, as indicated by the ICC, was excellent (Cicchetti, 1994) (3D FLAIR: mean ICC = 0.964; 2D FLAIR: mean ICC = 0.822). Detailed results on further metrics, and on segmented WMH volume can be found in **Supplementary Table 2**. In preparation for the optimization phase of the UBO Detector and BIANCA, as well as for the mandatory training dataset of BIANCA, the previously mentioned six 2D FLAIR images were manually segmented. **Figure 2** shows (left side) the mean WMH volumes resulting from the manual segmentations based on the 16 subjects per modality (T1w, 3D FLAIR, 2D FLAIR). No significant mean WMH volume differences were revealed between the three different manually segmented gold standards using the Friedman test with a Dunn Bonferroni post-hoc test (Holm correction). The Pearson’s product-moment correlation showed an almost perfect (Dancey & Reidy, 2017) linear association between all gold standards (mean *r* = 0.97, *p* < 0.001) (see **Figure 5** for the correlations between the gold standards but also between all three algorithms per input modality).

**Figure 2.**
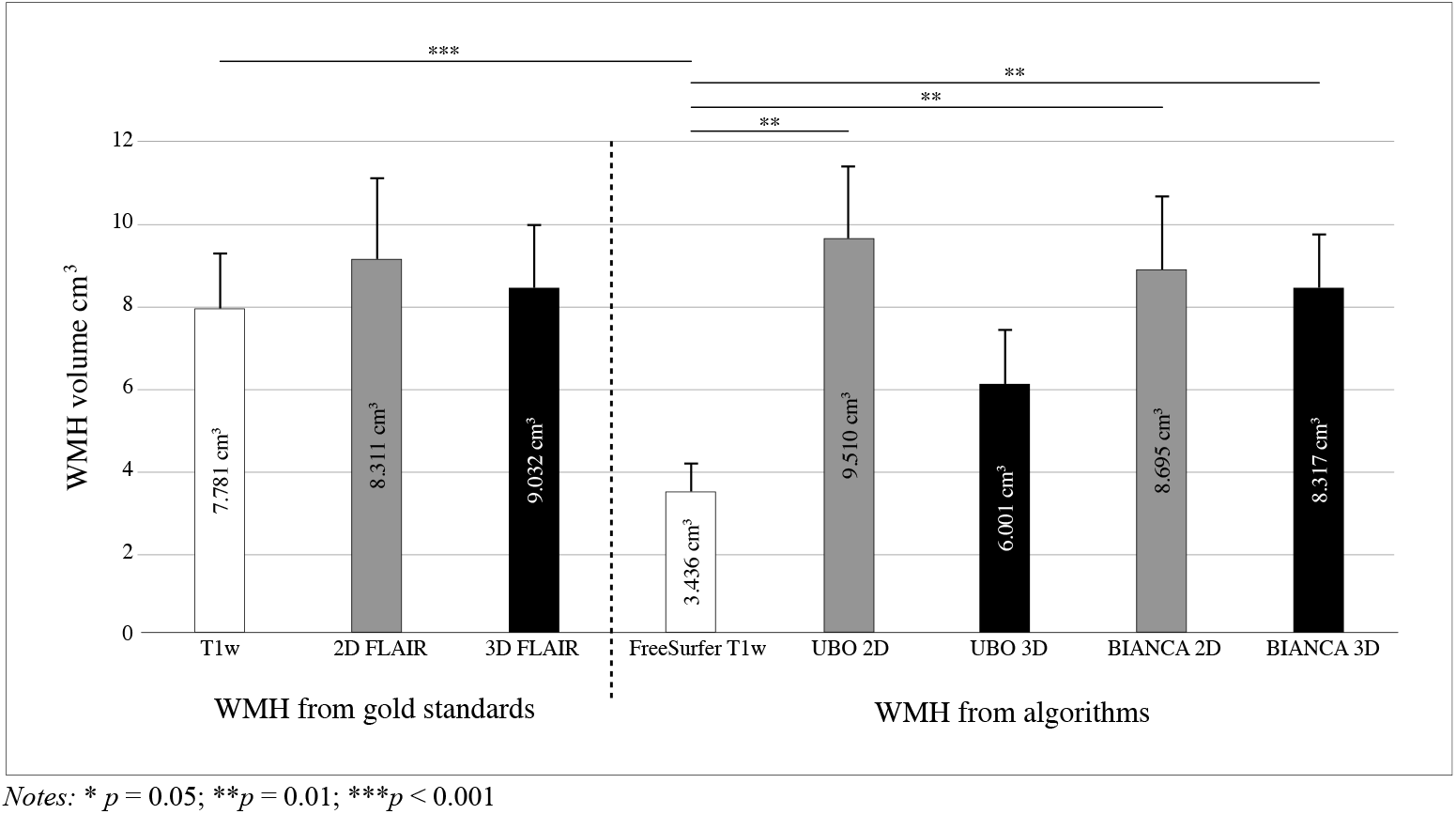
Mean WMH volume in cm^3^ with standard error of the mean (SEM) of the manually segmented gold standards on the left side of the figure, and the corresponding mean WMH volumes estimated by the automated algorithms on the right side of the figure.

### 2.6. Fazekas scale

The Fazekas scale is a widely used visual rating scale that provides information about the location (periventricular WMH (PVWMH) vs. deep WMH (DWMH)) and the severity of WMH lesions. It ranges from 0 to 3 for both locations, leading to a possible minimum score of 0, and a maximum score of 6 for total WMH.

In a first step, the three operators (O1, O2, O3) were specially trained by the neuroradiologist *S.K*. for several weeks on evaluating WMH with the Fazekas scale. *S.K*. was blinded to the demographics and neuropsychological data of the participants. 800 images were then visually rated using the Fazekas scale. Before the operators provided a final rating, they compared whether a given Fazekas score in the FLAIR images had the same score in the T1w images. We did not find differences in Fazekas scores between image modalities in any of the subjects. The ratings were carried out independently by all three operators, validated for the further procedure with statistical indicators that are described as follows. The inter-operator mean concordances across all four time points were determined with Kendall’s coefficient of concordance (Moslem, Ghorbanzadeh, Blaschke, & Duleba, 2019) by calculating it for total WMH, DWMH and PVWMH separately for each time point. Strong agreements according to Moslem et al. (2019) for total WMH (*W* = 0.864,*p* < 0.001), PVWMH (*W* = 0.828,*p* < 0.001), and DWMH (*W* = 0.842, *p* < 0.001) were found. Furthermore, mean inter-operator reliabilities were evaluated between the three operators for total WMH, PVWMH and DWMH across the four time points by using a weighted Cohen’s kappa (Cohen, 1968). Substantial to near-perfect reliabilities according to Landis and Koch (1977) were found (see **Supplementary Table 3**). The median score was calculated for each participant for each time point, split into total WMH, PVWMH and DWMH. The median Fazekas scores for all three operators were: total WMH = 3; PVWMH = 2; DWMH = 1. For more descriptive details see **Supplementary Table 4**.

### 2.7. Automated WMH segmentation

In order to obtain the best performance from each algorithm, we chose the settings that were considered the best by the original authors of UBO Detector and BIANCA. Thus, we did not intend to examine comparable settings (e.g., for the PVWMH), but rather to compare the outcome of the algorithms under optimal conditions.

#### 2.7.1. FreeSurfer

The FreeSurfer Image Analysis Suite (Fischl, 2012) uses a structural segmentation to identify regions in which WMH can occur, while regions in which WMH cannot occur are excluded (cortical and subcortical gray matter structures). The algorithm assigns a label to each voxel based on probabilistic local and intensity-related information that is automatically estimated from a built-in training dataset comprising 41 manually segmented images (surfer.nmr.mgh.harvard.edu/fswiki/AtlasSubjects) (Fischl et al., 2002). The algorithm differentiates between hypointensities in the white (WMH) and grey matter (non-WMH). The T1w images (subset 1; T1w images) were processed with FreeSurfer v6.0.1 as implemented in the FreeSurfer BIDS-App (Gorgolewski et al., 2017).

#### 2.7.2. UBO Detector

UBO Detector was applied to subset 2 (2D FLAIR + T1w) and subset 3 (3D FLAIR + T1w). UBO Detector calculates the probability of WMH by applying a classification model trained on ten subjects, manually segmented on 2D FLAIR images (built-in training dataset). A user-definable probability threshold generates a WMH map by segmenting the subregions including PVWMH, DWMH, lobar and arterial regions. As recommended by Jiang and colleagues (2018) we used a 12 mm threshold from the ventricular border to define the borders of PVWMH. The segmentation of the T1w images failed in five images, whereupon these subjects were excluded from the following procedures. Visual inspection of the data uncovered a segmentation error (eyeballs were marked as WMH), thus, this time point was excluded from further analysis leading to a total number of *N* = 756 subjects. To determine which settings are best suited for our subsets, we have evaluated the performance of four different settings proposed by Jiang et al. (2018), using the leave-one out cross-validation method. Since we had manually segmented the WMH for both 2D and 3D FLAIR images, we examined the performance of UBO Detector separately for each modality (2D FLAIR + T1w, 3D FLAIR + T1w). To do so, we calculated the accuracy metrics separately for the different settings, and checked which adjustments achieved the most optimal values (for further results see **Supplementary Table 5**). For 2D FLAIR images, UBO Detector worked most accurately with a threshold of 0.9 and a NN of *k* = 3. For the 3D FLAIR images, the best performance was achieved with a threshold of 0.7 and a NN of *k* = 5. For the subsequent calculations we used these optimized settings.

#### 2.7.3. BIANCA

BIANCA was applied to subset 2 (2D FLAIR + T1w) and subset 3 (3D FLAIR + T1w). For BIANCA a FLAIR training dataset is mandatory. As an output, BIANCA generates a probabilistic map of WMH for total WMH, PVWMH and DWMH. The 16 manually segmented gold standards derived once from 3D and once from 2D FLAIR images were used for the training datasets combined with the T1w images. As a first step, we compared the two thresholding options of BIANCA – best global threshold (0.99) vs. LOCATE to investigate which option provides the better segmentation quality for our subsets. To do so we used the leave-one out cross-validation method with the 16 gold standards per modality. We compared the outputted WMH with the DSC, DER, OER, H95, FP, TP, FPR, Sensitivity, ICC but also with the WMH volumes in cm^3^ by using the Wilcoxon-rank-sum-test. Because LOCATE did not perform better than the best global thresholding with 0.99 we used the latter for further analyses. For more information, and the detailed comparison analysis, see **Supplementary Analysis.**

To verify whether 16 gold standards for the training dataset were sufficient, we used the BIANCA *evaluation script*. BIANCA showed good results for both modalities (see **Supplementary Table 7**). For defining the PVWMH we adopted the 10 mm distance rule from the ventricles (DeCarli et al., 2005), which was also suggested by Griffanti et al. (2016). To reduce false positive voxels in the grey matter, and at the same time only localize WMH in the white matter, we applied a WM mask. For the BIANCA options we chose the ones that Griffanti et al. (2016) indicated as the best in terms of DSC and cluster-level false-positive ratio: MRI modality = FLAIR + T1w, spatial weighting = «1», patch = «no patch», location of training points = «noborder», number of training points = number of training points for WMH = 2000 and for non-WMH = 10000. For more details on the descriptions and the options, see Griffanti and colleagues (2016).

The preprocessing steps, applied before the BIANCA segmentation procedure, were performed with a nipype pipeline (v1.4.2; (Gorgolewski et al., 2011) as follows: Based on subject-specific template created by the anatomical workflows of fMRIprep (v1.0.5; (Esteban et al., 2019)), a WM-mask (FSL’s make_bianca_mask command) and a distancemap (distancemap command) were created. The distancemap was thresholded into periventricular and deep WM (cut-off = 10 mm). For each session, the two T1w images of a given time point were bias-corrected (ANTs v2.1.0; (Tustison et al., 2010)), brought to the template space, and averaged. FLAIR images were bias-corrected, and the template-space images were brought into FLAIR space using FLIRT (Jenkinson & Smith, 2001). To select the best threshold for the probabilistic output of BIANCA we first used the leave-one out cross-validation method to calculate the different validation metrics separately for the 2D and 3D FLAIR gold standard images (+ T1w images). The global threshold values 0.90, 0.95, 0.99 were applied, with the threshold of 0.99 for both FLAIR sequences proving to be the best fitting. For a more detailed overview see **Supplementary Table 6**.

### 2.8. Statistical analysis

According to the study questions outlined at the end of the introduction, we subdivided our analyses in two parts. In the first part, we assessed the accuracy of the segmentation methods by comparing the WMH estimates provided by the different algorithms with the corresponding gold standard. In the second part, we examined associations between the estimated WMH volumes, the Fazekas score and age.

#### Part 1 analyses

1a) First, we calculated the accuracies of the automatically extracted WMH segmentations by comparing them to the corresponding gold standard segmentations separately for the different algorithms using the above-mentioned accuracy metrics: DSC, DER, OER, H95, FPR, Sensitivity, and ICC (for results see **Table 4**). These accuracy metrics – except for the ICC – were compared using a Friedman test with a Dunn-Bonferroni post-hoc test (Holm correction). The ICC comparisons were interpreted as reliability measures according to Cicchetti (1994). For significant post-hoc results, Cohen’s *d* (Cohen, 1988) effect sizes were calculated.
1b) Secondly, we compared the automatically estimated WMH volumes amongst each other and with the volumes determined from the manually segmented gold standards. The results are summarized in **Figure 2**. For the statistical comparisons we used a Friedman test, because of a non-normal distribution, with a Dunn-Bonferroni post-hoc test (Holm correction) (Friedman, 1937).

**Table 4.** Summary of the accuracy metrics for the WMH using the three different modalities (T1w, 3D FLAIR, 2D FLAIR). DSC: Dice Similarity Coefficient, H95: Hausdorff distance for the 95th percentile, sensitivity, FPR: false positive ratio, DER: Detection Error Rate, OER: Outline Error Rate, ICC: Interclass Correlation Coefficient (ICC). Gray shadowed cells indicate significantly worse performance.

|  | FreeSurfer (T1w) |  | UBO (3D FLAIR) |  | UBO (2D FLAIR) |  | BIANCA (3D FLAIR) |  | BIANCA (2D FLAIR) |  |  |  |
| --- | --- | --- | --- | --- | --- | --- | --- | --- | --- | --- | --- | --- |
|  | Mean | 95% CI | Mean | 95% CI | Mean | 95% CI | Mean | 95% CI | Mean | 95% CI | *p*-value | Post-hoc<br>Dunn-Bonferroni (Holm correction) |
| **DSC** † | 0.434 | 0.375 – 0.493 | 0.500 | 0.435 – 0.567 | 0.531 | 0.471 – 0.591 | 0.602 | 0.531 – 0.672 | 0.561 | 0.498 – 0.624 | 0.001 | BIANCA 3D > FreeSurfer |
| **H95 (mm)** ‡ | 8.660 | 6.804 – 10.515 | 11.533 | 8.520 – 14.545 | 13.522 | 10.324 – 16.720 | 6.200 | 4.093 – 8.305 | 8.443 | 6.421 – 10.464 | < 0.001 | BIANCA 3D > UBO 3D, UBO 2D |
| **Sensitivity** † | 0.315 | 0.260 – 0.370 | 0.427 | 0.353 – 0.501 | 0.572 | 0.500 – 0.643 | 0.611 | 0.538 – 0.683 | 0.575 | 0.492 – 0.657 | < 0.001 | UBO 2D, BIANCA 3D, BIANCA 2D > FreeSurfer; BIANCA 3D > UBO 3D |
| **FPR** † | 4.62 $E^{-5}$ | 0.0000 – 0.0001 | 0.0002 | 0.0001 – 0.0002 | 0.0004 | 0.0003 – 0.0006 | 0.0003 | 0.0001 – 0.0004 | 0.0004 | 0.0002 – 0.0005 | < 0.001 | FreeSurfer < all others;<br>UBO 3D < UBO 2D |
| **DER** † | 0.200 | 0.146 – 0.253 | 0.313 | 0.228 – 0.398 | 0.327 | 0.231 – 0.423 | 0.162 | 0.113 – 0.210 | 0.215 | 0.143 – 0.287 | < 0.001 | BIANCA 3D < UBO 3D, UBO 2D |
| **OER** † | 0.932 | 0.839 – 1.025 | 0.687 | 0.629 – 0.746 | 0.611 | 0.540 – 0.683 | 0.635 | 0.538 – 0.733 | 0.664 | 0.586 – 0.742 | < 0.001 | all others < FreeSurfer |
|  | coeff | *p*-value | coeff | *p*-value | coeff | *p*-value | coeff | *p*-value | coeff | *p*-value |  |  |
| **ICC(3,1)** | 0.454 | 0.081 | 0.876 | < 0.001 | 0.927 | < 0.001 | 0.743 | < 0.05 | 0.859 | < 0.001 |  |  |

Notes: † Comparison by Friedman. ‡ A direct comparison is only allowed between the same modalities.

#### Part 2 analyses

2a) In order to examine how strongly the subjective Fazekas scores and the estimated WMH volumes correspond, we calculated the Spearman’s rho between these measures (**Table 5**). We examined these correlations across all time points but also separately for each time point since the sample sizes substantially decreased from time point to time point. For UBO Detector and BIANCA we analyzed PVWMH and DWMH volumes in addition to the total WMH volumes. The Spearman’s rho was used since the Fazekas scores are ordinally scaled.
2b) To estimate the influence of age on total WMH volumes, we applied linear mixed models (LMMs) with multiple measurements nested within individuals to satisfy the longitudinal nature of the used dataset. Total WMH volumes represented the dependent variables and entry age was entered as independent variable (see **Table 6**). We used the estimated total WMH volumes provided by the different segmentation methods, and calculated five LMMs. The WMH volumes and age were log-transformed and z-standardized. Relative effect sizes were calculated following Brysbaert and Stevens (2018) and according to Westfall, Kenny, and Judd (2014). The analyses were carried out in R (R Core Team, 2020) using the lme4 package (Bates, Mächler, Bolker, & Walker, 2015). Since we were interested in the linear associations, we reported the fixed effects (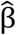 = standardized beta) of age at baseline for the different algorithms. Furthermore, the LMMs allows a comparison of the error variance. This measurement depends on the deviations from the individually estimated trajectories, of which – at least a part – can be considered as measurement errors of the algorithms.

**Table 5.**
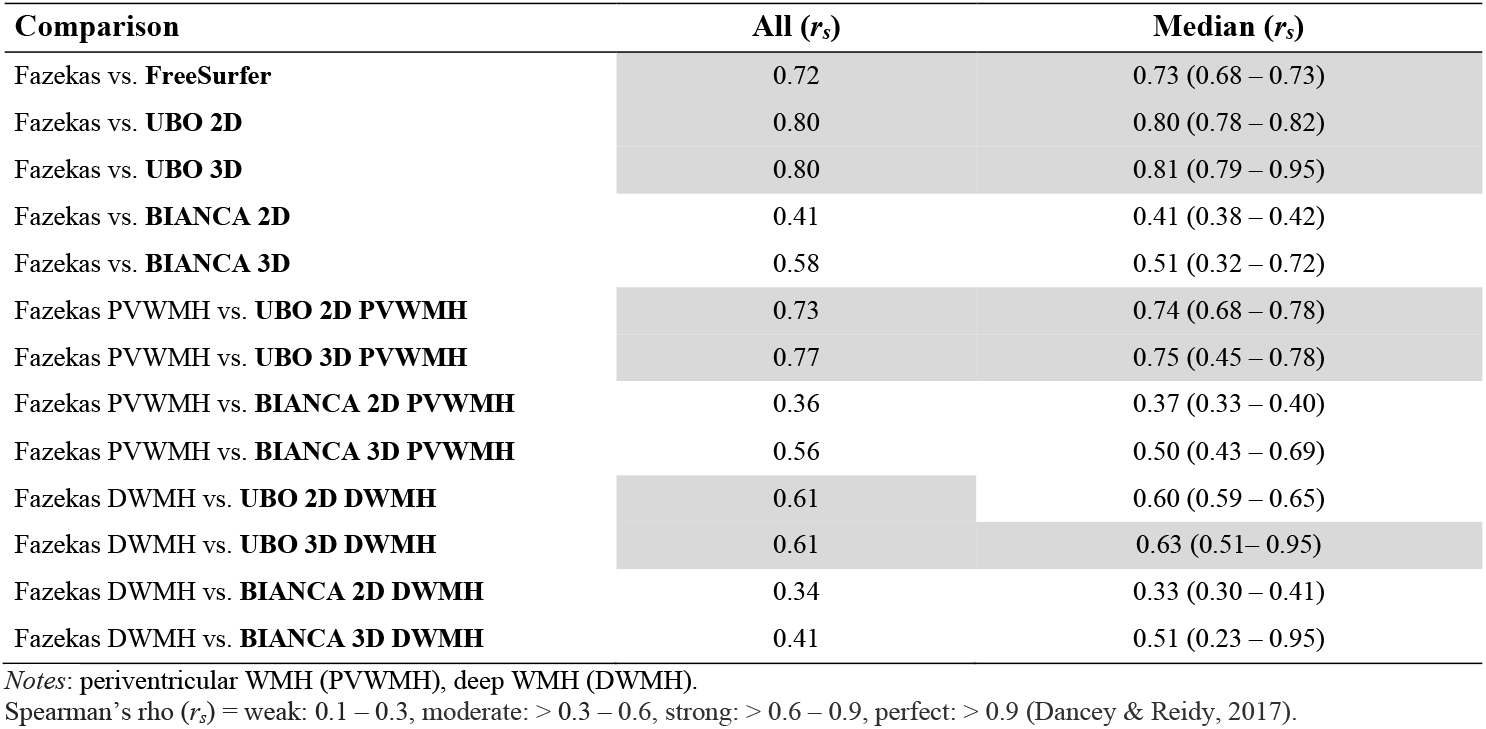
Fazekas scores versus WMH volumes. Spearman’s rho between the Fazekas scores and the different WMH volume measures. Shown are the correlations (r_s_) across the entire sample (All) and the median (Median) correlation across all four time points (minimum and maximum correlations are shown in brackets). Correlations r_s_ > 0.6 are highlighted by grey shading.

**Table 6.** Summary of the main effect of age on WMH volumes.

|  |  | **FreeSurfer T1w** | **UBO 2D** | **UBO 3D** | **BIANCA 2D** | **BIANCA 3D** |
| --- | --- | --- | --- | --- | --- | --- |
|  |  | ***N* = 800** | ***N* = 756** | ***N* = 166** | ***N* = 762** | ***N* = 166** |
| **Main effect**<br>**WMH ~ Age** | Age entry † | $\hat{\beta} = 0.455^{***}$ | $\hat{\beta} = 0.408^{***}$ | $\hat{\beta} = 0.306^{***}$ | $\hat{\beta} = 0.293^{***}$ | $\hat{\beta} = 0.339^{***}$ |
|  | CI | 0.347 – 0.564 | 0.295 – 0.520 | 0.127 – 0.485 | 0.183 – 0.403 | 0.173 – 0.506 |
|  | Cohen's *d* ‡ | 0.594 | 0.482 | 0.294 | 0.326 | 0.368 |
| **Residuals** | Estimates | 0.116 | 0.161 | 0.132 | 0.434 | 0.417 |
|  | CI | [0.109 – 0.123] | [0.152 – 0.172] | [0.109 – 0.163] | [0.409 – 462] | [0.344 – 0.520] |

*Notes:* 95% confidence interval (CI).
† $\hat{\beta}$ = (standardized beta) fixed effect (slope). The WMH volumes are log-transformed and z-standardized. Chronological age is z-standardized. ‡ Cohen's *d*: small effect size: $\geq 0.2$ –, medium effect size: 0.5, large effect size: 0.8 (Cohen, 1988).
\*\*\* $p < 0.001$

Finally, we ran post-hoc calculations based on our observations of strong discrepancies between WMH volumes of consecutive time points in the BIANCA output, which we refer to as «outlier intervals» in the following. To further investigate the within-subject variability of WMH volumes, we determined the percentage and number of «outlier intervals between two measurement points» and also of «subjects with outlier in longitudinal data» based on the mean percentages of WMH volume increases (mean + 1SD) and decreases (mean - 1SD), separately for possible time intervals (1-, 2-, 3-, 4-year intervals) to calculate a «range of tolerance» and exclude outlier data points. The outlier data points were further classified for low, medium, and high WMH load (based on the Fazekas scale) in order to identify a specific pattern. If the number of «outlier intervals between two measurement points» exceeded the number of «subjects with outlier in longitudinal data», this indicated peaks or even several outlier data points within a single person – and would be an indication of a zigzag pattern over time. For a more detailed description see ***Post-hoc outlier analysis*.**

For all statistical analyses appropriate R packages were applied. Where possible, differences were also expressed as Cohen’s *d* (Cohen, 1988). A *d* of 0.2 is considered as small, a *d* of 0.5 as medium, and a *d* of 0.8 as large effect size (Cohen, 1988). Spearman’s rho are classified according to Dancey and Reidy (2017) (*r_s_* > 0.1 – 0.3 as weak, *r_s_* > 0.3 – 0.6 as moderate, *r_s_* > 0.6 – 0.9 as strong, and *r_s_* > 1 as perfect). For post-hoc tests in the context of the Friedman tests we performed adjusted Bonferroni paired t-tests. The reliability estimates on the basis of the ICCs were classified according to Cicchetti (1994) (ICC = fair: > 0.4 – 0.6, good: > 0.6 – 0.75, excellent > 0.75).

### 2.9. Computer equipment

All WMH extractions were undertaken on a Supermicro X8QB6 workstation with 4 × Intel Xeon E57-4860 CPU (4 x10 cores, 2.27 GHz) and 256 GB RAM. The computing host was a KVM virtualized guest instance with Ubuntu 18.04.4 LTS with 32 x Intel Xeon E7-4860 CPU (2.27 GHz) and 92 GB RAM.

## 3. Results

### 1a) Accuracies of the WMH segmentation methods

**Table 4** summarizes the accuracies of the WMH segmentation algorithms based on the comparison between the algorithm outputs and the corresponding gold standards.

Generally, the accuracies are at least fairly good. Comparing the accuracy measures – using the Friedman test – revealed that FreeSurfer underperforms in important accuracy measures like DSC, sensitivity, and OER compared to the other algorithms. Regarding the ICC (range: 0.45 to 0.93) the reliability according to Cicchetti (1994) revealed a fair reliability for FreeSurfer, a good reliability for BIANCA 3D, and excellent reliabilities for UBO Detector 2D and 3D FLAIR, and BIANCA 2D FLAIR. In summary, FreeSurfer showed the weakest performance in terms of segmentation accuracy, and BIANCA 3D FLAIR was the most accurate. Differences between UBO Detector and BIANCA were generally very small and rarely reached significance, especially when applying the algorithms to 2D FLAIR images.

The DSC between gold standard and the respective algorithm’s output divided into low, middle and high WMH load is shown in **Supplementary Table 9.**

### 1b) Relationship between the gold standard WMH volumes and automatically estimated WMH volumes

**Figure 2** depicts the WMH volumes of the different gold standards and the automatically estimated WMH volumes of the same brains. These WMH volumes were subjected to a Friedman test, which revealed a significant result (Chi-Square(7) = 56.375,*p* < .001, *n* = 16). The pairwise comparisons revealed no significant differences between the gold standards. Comparing the automatically estimated WMH volumes with their corresponding gold standard WMH volumes revealed a substantial difference between those from the gold standard T1w and those estimated by FreeSurfer (*p* < 0.001). The comparisons among the algorithm outputs showed a significantly lower WMH volume estimate of FreeSurfer compared to the estimates provided by BIANCA 2D and 3D (*p* = 0.01) as well as UBO 2D (*p* = 0.01). All further pairwise comparisons (e. g., 2D FLAIR gold standard vs. UBO 2D, 2D FLAIR gold standard vs. BIANCA 2D etc.) showed no significant differences.

### 2a) Relationship between the subjective Fazekas scores and the estimated WMH volumes

**Table 5** summarizes the relationship between the estimated WMH volumes and the Fazekas scores in terms of Spearman’s rho (*r_s_*). Seven correlations out of 13 for the entire sample are strong (> 0.6). The remaining correlations for this sample are moderately strong. Considering the median correlations across the four time points revealed very similar results. **Table 5** also demonstrates the range of the correlations among the time points. The spaghetti plots for total WMH volume and Fazekas scores are depicted in **Figure 3A.**

**Figure 3.**
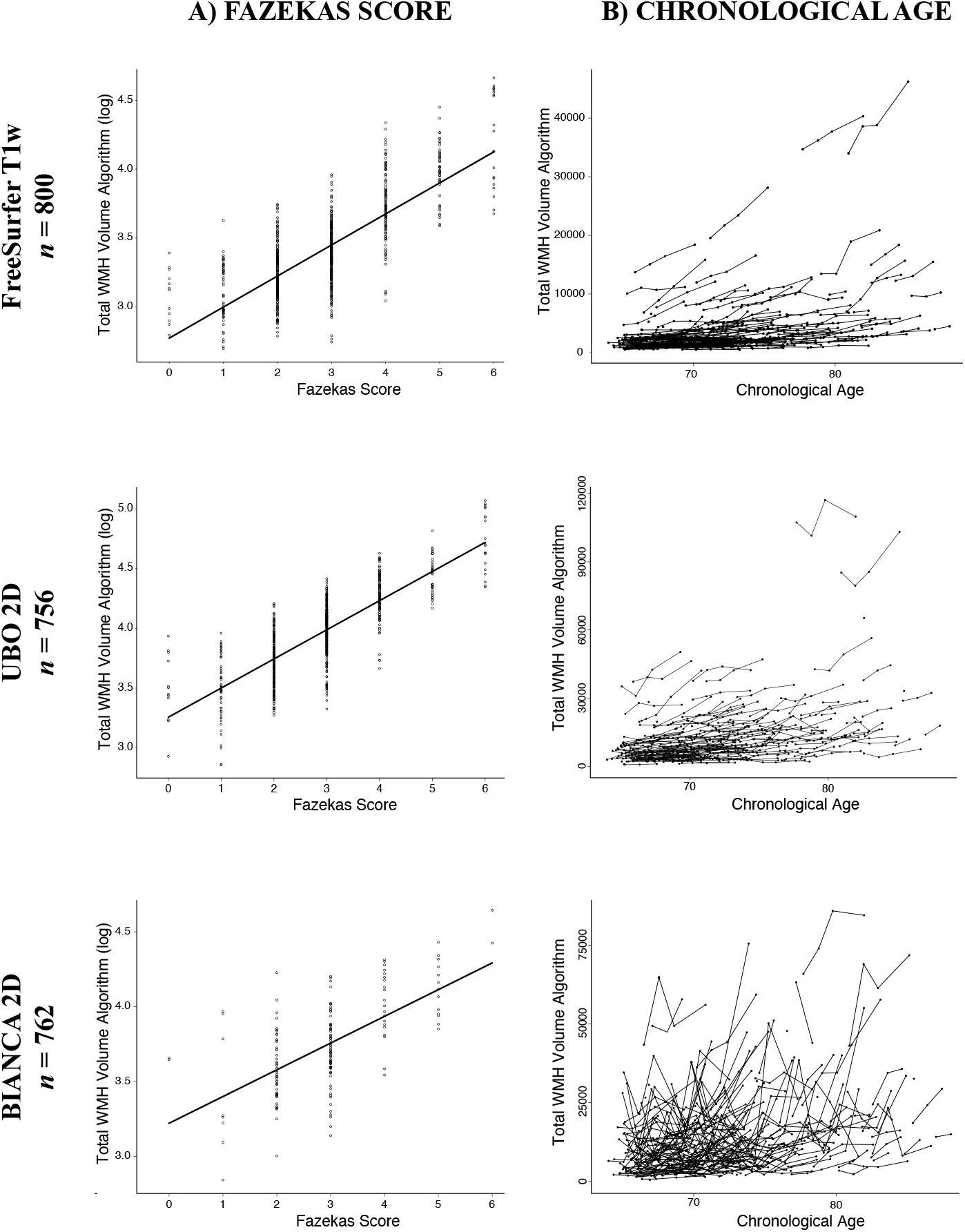
Validation of the three algorithms with the subsets FreeSurfer T1w, UBO 2D, BIANCA 2D. Scatter plot of the log-transformed total WMH volume distribution according to Fazekas score (A), and spaghetti plot of total WMH volume and chronological age (B).

### 2b) Relationship between the estimated WMH volumes with age and the longitudinal time course

**Table 6** summarizes the results of the LMM regression analysis with the WMH volumes as dependent variables and chronological age as independent variable. As one can see in this table, chronological age is moderately associated with all total WMH volume measures. The corresponding effect sizes for this association range between 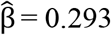 (BIANCA 2D) and 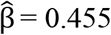 (FreeSurfer). The spaghetti plots for total WMH volumes and chronological age are shown in **Figure 3B**.

### Post-hoc outlier analysis

The results described above indicate that BIANCA demonstrates increased variability. A close inspection of this variability revealed that the WMH segmentations masks contained erroneous voxels, particularly in the following areas: semi-oval center, orbitofrontal cortex (orbital gyrus, gyrus rectus, above the putamen), and occipital lobe below the ventricles. Although recent studies indicate some WMH variability (Shi & Wardlaw, 2016), we assumed that these massive peaks are driven by outliers. Since our knowledge of WMH volume changes in healthy older adults is insufficient (Shi & Wardlaw, 2016) and has inconsistent findings (Ramirez, McNeely, Berezuk, Gao, & Black, 2016), we identified outliers in WMH volume increases and decreases, based on average WMH volume changes in our sample. Please find a detailed description of our approach in **Supplement** «**Detailed description of post-hoc outlier analysis»**.

BIANCA 2D and 3D clearly showed more outlier intervals than the other algorithms (BIANCA 2D: *n* = 161 outlier intervals, 30.32%; BIANCA 3D: *n* = 8, 16.67%), see **Table 7**. Furthermore, BIANCA 2D/ 3D was the only algorithm with more outliers than persons which is reflective of the zigzag patterns within the subject’s trajectories. Please see **Supplementary Table 10** for more results on WMH volume changes in percent. There was no clear association between the number of outliers and lesion load; see **Supplementary Table 11.**

## 4. Discussion

In this study we assessed the performance of three freely available and automated algorithms for WMH segmentation: FreeSurfer, UBO Detector, and BIANCA. To do so, we applied the algorithms to a large longitudinal dataset comprising T1w, 2D FLAIR and 3D FLAIR images from cognitively healthy, older adults with a low WMH load. We discovered that all algorithms have certain strengths and limitations. FreeSurfer showed deficiencies particularly with respect to segmentation accuracy (i.e., DSC) and clearly underestimated the WMH volumes. We therefore argue that it cannot be considered as a valid substitution for manually segmented WMH. BIANCA and UBO Detector showed a higher segmentation accuracy compared to FreeSurfer. When using 3D FLAIR + T1w images as input, BIANCA performed significantly better than UBO Detector regarding the accuracy metrics DER and H95. However, for BIANCA we identified a significant number of outliers in the individual trajectories of the WMH volume estimates. UBO Detector – as a fully automated algorithm that works without a training dataset – showed the best cost/benefit ratio in terms of processing time and segmentation performance in our study. Although there is room for optimization regarding segmentation accuracy, it distinguished itself through its excellent volumetric agreement with the manually segmented WMH in both FLAIR modalities (as reflected by the ICCs) and its high correlations with the Fazekas scores. In addition, it proved to be a robust estimator of WMH volumes over time.

### 4.1. Evaluation of the algorithms

#### FreeSurfer

The total WMH volumes provided by FreeSurfer showed a strong correlation with the Fazekas scores and a moderate association with age. FreeSurfer showed no within-person outlier WMH volume estimations. The biggest constraint is the fundamental underestimation of WMH volume compared to the corresponding T1w gold standard, which compromises the validity of its output. The underestimation can be attributed to the fact that WMH often appear isointense in T1w sequences and are therefore not detected (Wardlaw et al., 2013). Furthermore, the lower contrast of the DWMH compared to the PVWMH, which is due to the lower water content in the DWMH as a result of the longer distance to the ventricles, might contribute to the WMH volume underestimation. FreeSurfer often omitted DWMH, a finding also reported by Olsson et al. (2013). In addition, our analyses showed that FreeSurfers’ underestimation of the WMH volume was even more pronounced in high WMH load images (see Bland-Altmann Plot in **Figure 4**, panel C). The same bias was shown by Olsson et al. (2013) when comparing the WMH volumes of the semi-manually segmented WMH (2D FLAIR) and the FreeSurfer (T1w) output. One reason for the underestimation in subjects with high WMH loads might be the fact that FreeSurfer segments the WMH as grey matter (e.g., for the bilateral caudate) (Dadar, Potvin, Camicioli, & Duchesne, 2020) which could also explain FreeSurfer’s low false positive ratio. The spatial overlap performance of FreeSurfer in our study is comparable to the findings in the validation study reported by Samaille and colleagues (2012) with a cohort of mild cognitive impairment and CADASIL patients. Although other studies reported higher volumetric agreement between FreeSurfer’s output and manual segmentation (Ajilore et al., 2014; Smith et al., 2011), the WMH volume difference between T1w and both FLAIR modalities is in line with the STRIVE, stating that FLAIR images tend to be more sensitive to WMH and therefore are considered more suitable for WMH detection than T1w images (Dadar et al., 2018; Wardlaw et al., 2013). However, to our knowledge, most previous studies do not indicate whether they used a fully manually segmented gold standard or a gold standard generated by a semi-automated method. Further, the study samples have been small, there is very little information on commonly used accuracy metrics like DSC, DER, OER etc., and FreeSurfer’s WMH algorithm has not been applied to longitudinal data. Moreover, previous studies have not compared FreeSurfer’s WMH volumes with manual segmentations on T1w structural images or with visual rating scales such as the Fazekas scale. In our study, FreeSurfer’s WMH volumes strongly correlated with the Fazekas scores and showed reliable WMH volume estimations across time points. However, FreeSurfer cannot be considered as a valid substitute for manual WMH segmentation on this dataset due to the weak outcomes in the accuracy metrics (DSC, OER, ICC), and especially due to its massive WMH volume underestimation. Nevertheless, because of the valid and reliable outcomes with the Fazekas scale, FreeSurfer is suitable for use in clinical practice, as long as its values are not interpreted as absolute values.

**Figure 4.**
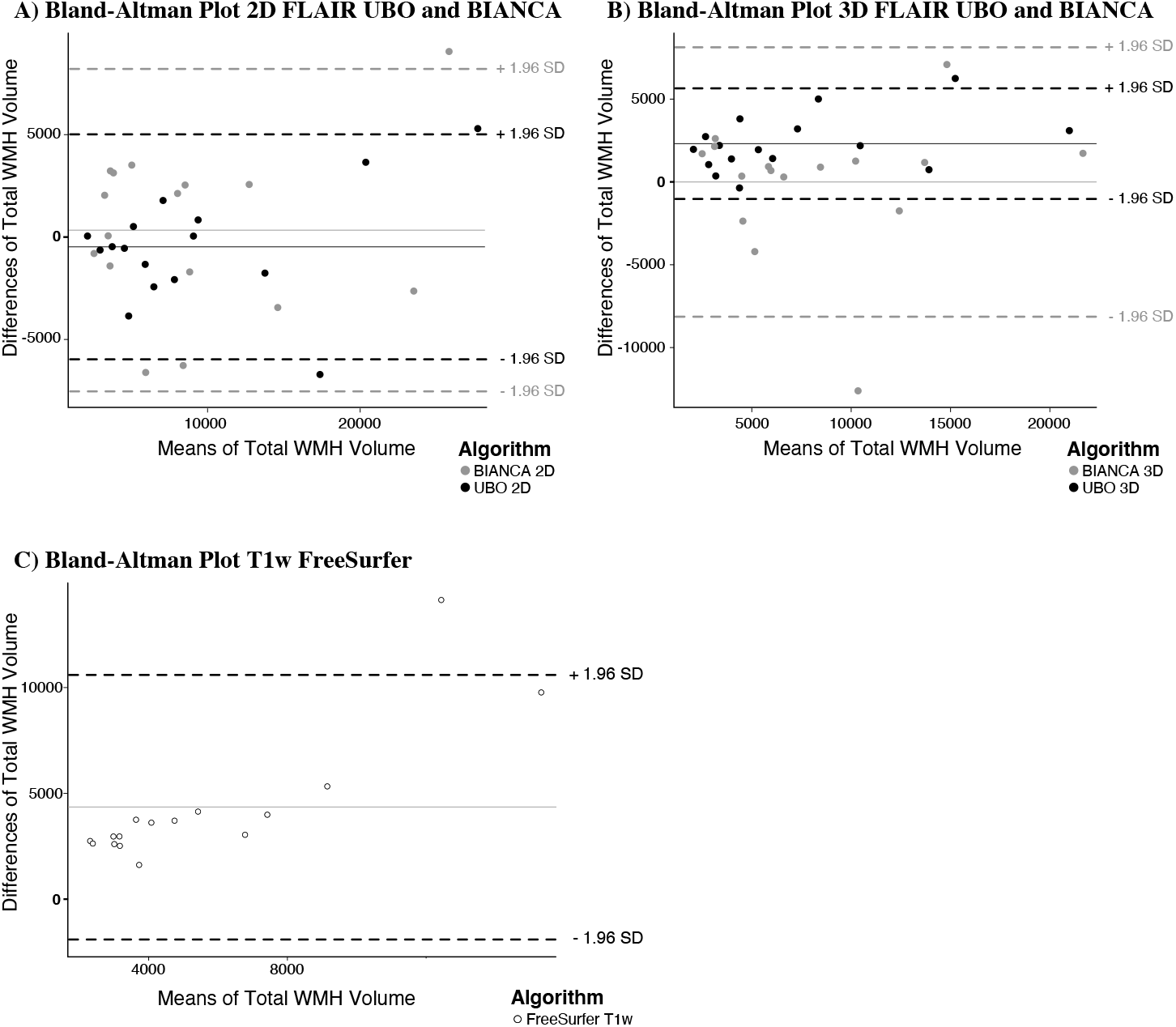
Bland-Altman (Bland & Altman, 1986) plots for WMH volume for the different algorithms (total WMH volume gold standard minus total WMH volume algorithm in mm^3^). The x-axes contain mean WMH volumes, the y-axes contain absolute differences of WMH volumes: S (x, y) = (gold standard WMH volumes (S1) + algorithm WMH volumes (S2)/2, S1 – S2).

#### UBO Detector

The total WMH volumes estimated by UBO Detector with the 2D FLAIR + T1w and 3D FLAIR +T1w inputs were strongly correlated with the Fazekas scores and showed significant relations with age. PVWMH and DWMH volumes with both FLAIR input modalities showed strong, and moderate correlation with Fazekas scores, respectively. This is consistent with the article by the developers of UBO Detector (Jiang et al., 2018), which reported significant relations between UBO Detector-derived PVWMH and DWMH volumes and the Fazekas scores. The results of their volumetric agreements – calculated with ICCs – were similar to ours, especially for UBO 2D, but we were not able to replicate the high values they obtained in sensitivity and overlap measurements (DSC, DER, and OER). Our DSC and ICC values were similar to those of the very recent cross-sectional study of Vanderbecq et al. (2020). The discrepancies between the results of the developers’ study and ours could be due to the fact that the built-in training dataset of UBO Detector is based on 2D FLAIR images, which may cause differences in WMH segmentation performance depending on the image input modality (2D vs. 3D FLAIR). To our knowledge, our study is the first study, to validate UBO Detector with 3D FLAIR images cross-sectionally and longitudinally. Indeed, our analysis indicated that the WMH volumes estimated by UBO Detector depend on the modality of the FLAIR input. Volumes extracted with UBO 2D tended to be more similar to the WMH volume of the gold standard with the same modality, while volumes extracted with UBO 3D tended to underestimate the WMH volumes of the respective gold standard (see **Figure 2**, and Bland-Altmann Plot in **Figure 4**, panel A and B). For several reasons UBO Detector’s longitudinal pipeline was not used in our study. First, UBO Detector requires an equal number of sessions for all subjects, which would have resulted in a marked reduction of our sample size. Secondly, it registers all sessions to the first time point, an approach that has been shown to lead to biased registration (Reuter, Schmansky, Rosas, & Fischl, 2012). Lastly, comparing the two pipelines, Jiang et al. (2018) did not find significant differences regarding the extracted WMH volumes. To date, we have not found any other study that validated UBO Detector or compared it with other WMH extraction methods using a longitudinal dataset with different MR modalities.

#### BIANCA

To enable a direct comparison with the accuracy metrics used in the original BIANCA study (Griffanti et al., 2016) we additionally ran BIANCA’s *evaluation script* (**Supplementary Table 7**). Our overall results for the different accuracy metrics corresponded more to those of their vascular cohort than to those of their neurodegenerative cohort. We received quite similar results for BIANCA 2D with respect to the correlations between the estimated WMH volumes and our manually segmented WMH volumes. However, we were not able to replicate the high ICCs or the high associations they obtained between the WMH volumes and the Fazekas scores in both cohorts, either for BIANCA 2D or for BIANCA 3D. BIANCA 2D showed only a moderate correlation with the Fazekas scale – although we used a customized training dataset. With respect to age and WMH volumes, the small associations we obtained were similar to those reported by Griffanti et al. (2016) for their neurodegenerative cohort. We suspect that this might be due to the outlier segmentations in our BIANCA outputs. While the effect of outlier WMH segmentations was not detected in the smaller cross-sectional analyses, it was uncovered because of massive WMH volume fluctuations in the within-subject trajectories. When the developers of UBO Detector compared their algorithm with BIANCA, they noticed that BIANCA tended to overestimate the WMH in «milky» regions, whereas the sensitivity for WMH detection was higher in BIANCA than in UBO Detector, which is in line with our findings.

BIANCA features LOCATE as a method to determine spatially adaptive thresholds in different regions in the lesion probability map. Sundaresan et al. (2018) showed that LOCATE is beneficial when the BIANCA algorithm is trained with dataset-specific images or when the training dataset was acquired with the same sequence and the same scanner. For the group of healthy controls, they achieved similar visual outputs with LOCATE as compared to those with global thresholding. However, since no manually segmented gold standard was available for the healthy controls in their study, a quantitative comparison between manually segmented WMH and LOCATE’s WMH is lacking. In our analysis, LOCATE, as compared to BIANCA’s global thresholding did perform significantly worse at processing images with a low WMH load (see **Supplementary Analysis**). LOCATE undoubtedly had more true positives, resulting in significantly higher sensitivity, but this came at the cost of a 3-fold higher false positive rate (see **Supplementary Table 12**). Hence, all other metrics (DSC, OER, DER, H95 and FPR) showed worse outcomes for LOCATE compared to BIANCA’s global thresholding in our dataset. In addition, the WMH volumes LOCATE provided deviated significantly from the WMH volumes of the gold standards due to the high number of false positives in LOCATE. Ling and colleagues (2018) validated BIANCA with different input modalities (single FLAIR or FLAIR + T1w), in a cohort of patients with CADASIL using a semi-manually generated gold standard of ten subjects per modality. In their dataset, which contained an extremely high WMH load, they received higher DSC metrics for 2D and 3D images compared to our results, while the volumetric agreement was very similar to ours with the global threshold method. In the study by Vanderbecq et al. (2020), the ICC determined with their «clinical routine data set» (patients who were referred for assessment of cognitive impairment) with a 3D FLAIR + T1w image input was comparable to ours with the 2D FLAIR + T1w input, whereas the ICC determined with their «research data set» (ADNI dataset; comprising mainly Alzheimer’s and amnestic mild cognitive impairment patients) with a merged dataset of 2D and 3D FLAIR + T1w images as input, was lower compared to ours. Ling et al. (2018) found that BIANCA tended to overestimate the WMH volumes in subjects with a low WMH load and underestimate it in subjects with a high WMH load. According to them, in a group of healthy elderly people with a low WMH exposure, such a bias would be unlikely to be identified. In both BIANCA 2D and 3D we did not detect systematic biases but revealed one clear underestimation in the subject with the highest WMH load in the 2D FLAIR images, and one pronounced overestimation in a subject with medium WMH load in the 3D FLAIR images (see Bland-Altman Plot **Figure 4**, panel A and B). With a similar approach, using the absolute mean WMH volume differences to the gold standards, we were able to show that the mean WMH volume differences of the WMH volumes of BIANCA are the results of random averaging over inaccurately estimated WMH volumes (see **Supplementary Table 8**). Given that the focus of our study was to compare different algorithms in terms of costs and benefits, we did not test other settings for BIANCA but adhered to the default settings suggested in the original BIANCA validation. To our knowledge, BIANCA and LOCATE have not yet been validated with a longitudinal dataset.

### 4.2. Comparison of the algorithms with each other

The quality assessment of the algorithms is critically based on fully manually segmented WMH (gold standards). In order to prove construct validity, 16 gold standards per modality (T1w, 2D FLAIR, 3D FLAIR) were correlated amongst each other and with the respective WMH volume outcomes of the algorithms (**see Figure 5**).

**Figure 5.**
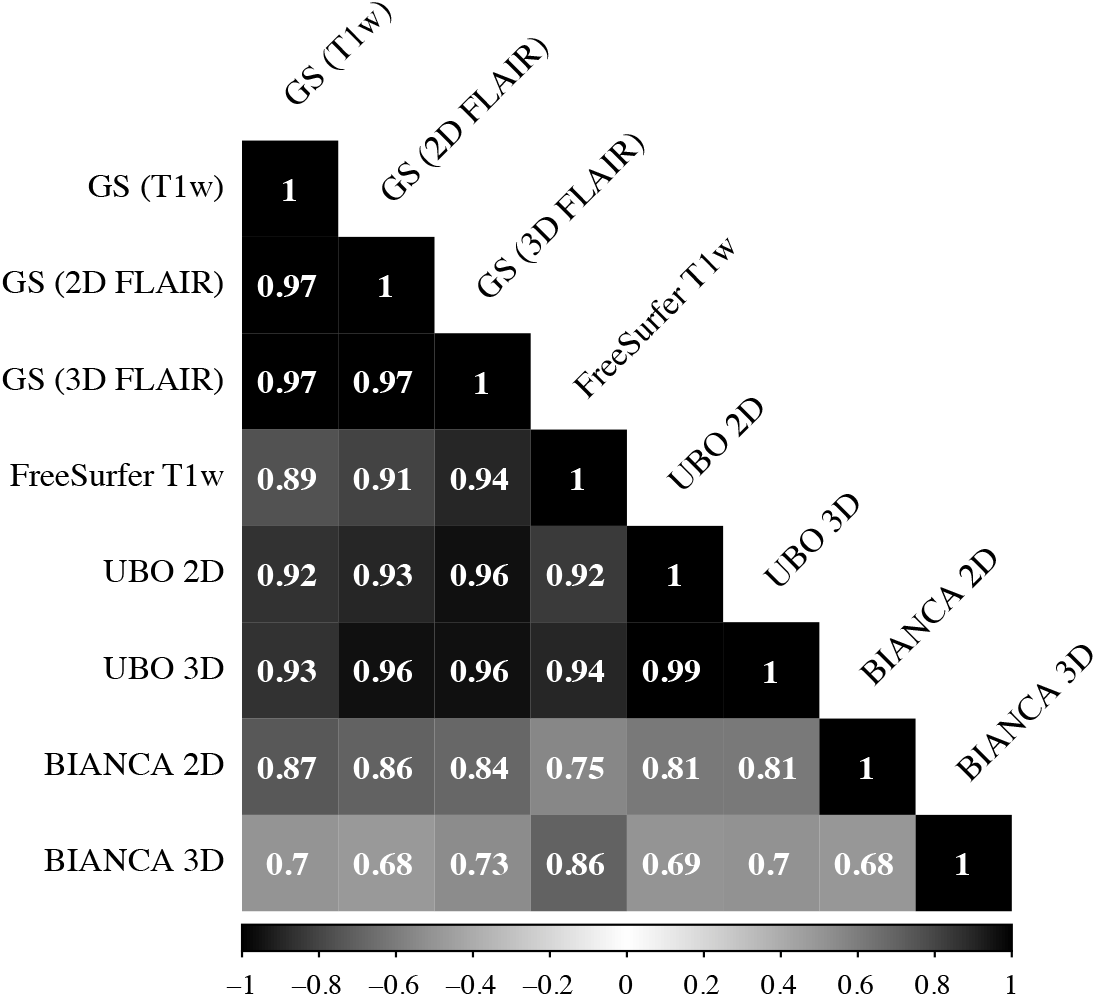
Correlation matrix of WMH volume estimation within the different gold standards (GS)(T1w), GS (2D FLAIR), GS (3D FLAIR), and between the gold standards and the different algorithms outputs (FreeSurfer T1w, UBO 2D, BIANCA 2D, UBO 3D, BIANCA 3D. Result of the correlation of the gold standards (all combinations: r = 0.97, p < 0.05).

The strong correlations (mean *r* = 0.97, *p* < 0.05) we identified between the WMH volumes of the gold standards indicate a very high validity for our gold standards. When evaluating association of the volumes provided by the different methods and the volumes of their respective gold standard, the highest correlations were found for UBO Detector 3D, followed by UBO Detector 2D, FreeSurfer T1w, BIANCA 2D, and BIANCA 3D. This correlation pattern is interesting and partly unexpected, especially when considering that BIANCA was the only algorithm that was fed with a customized training dataset for every modality (based on 2D and 3D FLAIR images). In line with this, very high associations were found between the WMH volumes estimates of UBO Detector 2D and 3D FLAIR, while the corresponding correlation was clearly smaller for BIANCA. Our results are consistent with the recent study of Vanderbecq et al. (2020), who also reported superior WMH segmentation accuracy of UBO Detector compared to BIANCA.

Notably, the WMH volumes of FreeSurfer correlated very highly with both UBO Detector outputs but less strongly with volumes estimates provided by BIANCA, which may be due to the outlier WMH segmentations BIANCA produced. On the other hand, the WMH volumes extracted by FreeSurfer were generally smaller than the outputs of UBO Detector and BIANCA. We would like to emphasize that FreeSurfer is the only algorithm that even underestimated its «own» gold standard. This WMH volume underestimation also affected the accuracy metrics (DSC, Sensitivity, OER and ICC), which were significantly worse for FreeSurfer compared to the other algorithms. Regarding H95 and DER, BIANCA 3D FLAIR performed best. On the flip side, BIANCA showed the weakest correlation with the Fazekas scores, and the larges residual variances in the LMMs. In line with the latter, BIANCA 2D and 3D FLAIR had the highest number of outlier WMH volumes compared to the other algorithms, which can be clearly detected in the within-subject trajectories. Some studies report annual percentage increases in WMH volumes in the range between 12.5% and 14.4% in subjects with early confluent lesions, and 17.3% and 25.0% in subjects with confluent abnormalities (Duering et al., 2013; Ramirez et al., 2016; Sachdev, Wen, Chen, & Brodaty, 2007; R. Schmidt et al., 2003; van Dijk et al., 2008). Ramirez et al. (2016) summarized the progression rates of WMH volumes in serial MRI studies in their Table 2, showing a wide variability of ranges. In our study, the annual WMH volume increases based on BIANCA’s estimations (e.g., 31.85% ± 76.5% (M±1SD) for BIANCA 2D) were clearly higher than the changes reported in the literature and they were also higher than the changes detected with UBO Detector and FreeSurfer in this study (see **Supplementary Table 10**), which is likely also related to the outlier segmentations that influenced segmentation’s reliability. Having a closer look on the segmentation variability of the algorithms by means of the Bland-Altman plots, which illustrate data of the subsample (*N* = 16) used for the manual segmentation (**see Figure 4**, panel A and B), we observed that the limits of agreement are wider in BIANCA than in UBO Detector. Moreover, the 2D and 3D FLAIR image plots show strong outliers (under- and overestimations). Interestingly, in this subsample the single deviations in BIANCA’s output seem to cancel each other out and result in a mean WMH volume that is very similar to the gold standard’s WMH volume (see **Supplementary Table 8**). From analyzing the validation of the algorithms, we can conclude that with our subsets UBO 2D /3D and FreeSurfer T1w, as compared to BIANCA 2D/3D, performed more robust and also more consistently across time. Future research needs to evaluate if the segmentation errors BIANCA produced with our longitudinal dataset also occur in the context of other datasets.

One general problem in the context of automated WMH lesion segmentation using FLAIR images is the incorrect inclusion of the septum pellucidum, the area separating the two lateral ventricles, in the output masks. This area appears hyperintense on FLAIR sequences, and therefore, looks very similar to WMH. When erroneously detected as WMH, the septum pellucidum enters the output volumes as a false positive region, which leads to an overestimation of the WMH volume. The UBO Detector developers also segmented the septum pellucidum in their gold standard (see their Supplementary Figure 1b). Since they already fed their algorithm with this false positive information, it was to be expected that UBO Detector would also segment the septum pellucidum in our data, which may have caused the worse DER compared to FreeSurfer and BIANCA.

### 4.3. Strength and limitations

The main strengths of this study are the validation and comparison of three freely available algorithms using a large longitudinal dataset of cognitively healthy adults. Importantly, we used fully manually segmented gold standards in all three planes (sagittal, coronal, axial) and for three different MR modalities (T1w, 2D FLAIR, 3D FLAIR) by multiple operators, who reached excellent inter-operator agreements. Besides the manual segmentations, our study features Fazekas scores for the whole dataset, which were used to cross-validate the WMH volumes provided by the segmentation algorithms. One limitation of this study is that we applied the algorithms to only one sample and that this sample was homogeneous with respect to its low lesion load. Future studies should determine how well these results generalize to other studies, scanners, sequences, and heterogeneous datasets including clinical populations.

### 4.4. Usability of the algorithms

Given that the algorithms for WMH extraction are usually not implemented by trained programmers, usability is an important issue to mention in the context of this work.

FreeSurfer has not been specifically programmed for WMH detection, but is a tool for extensive analysis of brain imaging data. Because of all the other parameters FreeSurfer outputs besides WMH volume, the processing time is very long (many hours per session). The FreeSurfer output comprises the total WMH volume and the total non-WMH volumes (grey matter).

UBO Detector has been specifically programmed for WMH detection, and has been trained with a «built-in» training dataset. Theoretically, it is possible to train the algorithm using a previously manually segmented gold standard. However, this procedure only works within the Graphic User Interface (GUI) in DARTEL space, and is very time-consuming. The output from UBO Detector is well-structured and contains among others the WMH volume and the number of clusters for total WMH, PVWMH, DWMH as well as WMH volumes per cerebral lobe. For subset 2, the whole WMH extraction process (incl. pre- and postprocessing) took approx. 14 min per brain with the computing environment specified in the methods section. For subset 3, the WMH extraction took approx. 32 min per brain.

BIANCA is a tool integrated in FSL (FMRIB’s Software Library) with no need of any other program. It is very flexible in terms of MRI input modalities that can be used and offers many different options for optimization. The output of BIANCA comprises the total WMH. If required, a distance from the ventricles can be selected to PVWMH and DWMH. In a longitudinal study with many subjects and time points, or also in a study with a big sample size, the aggregation of the algorithm output files seemed to be very time-consuming because of the many output files. Our preprocessing steps for BIANCA took about 2:40 h per subject for the preparation of the templates and about 1:10 h per session for the preparation of the T1w, 2D and 3D FLAIR images. After preprocessing, BIANCA required about 1:20 h per session for setting the threshold for both FLAIR images, and the WMH segmentation took about 4 min and 8 min per session for the 2D FLAIR and 3D FLAIR images, respectively.

## 5. Conclusions

The main aim of the current study has been the comparison and validation of three freely available algorithms for automated WMH segmentation using a large longitudinal dataset of cognitively healthy adults with a relatively low WMH load. Our results indicate that FreeSurfer underestimates the total WMH volumes significantly and misses some DWMH completely. Therefore, this algorithm seems not suitable for research specifically focusing on WMH and its associated pathologies. However, given the high associations with the Fazekas scores and its longitudinal robustness, FreeSurfer’s suitability for clinical practice could be further explored in future studies. BIANCA performed largely well with respect to the accuracy metrics. However, many outlier segmentations were identified when the algorithm was applied to the longitudinal dataset, which likely contributed to the lower correlations of BIANCAs WMH volume estimations with the volume estimations of the other algorithms as well as with the Fazekas scores and age. UBO Detector, as a completely automated algorithm, has scored best regarding the costs and benefits due to its fully generalizable performance. Although UBO Detector performed very well in this study, improvements in accuracy metrics, such as DER and H95, would be desirable for it to be considered as a true replacement for manual segmentation of WMH. In general, this study confirms the importance of validating the algorithms based on longitudinal datasets – especially in the studies with large samples, where it is not feasible to visually check and verify every single image and its WMH segmentation.

## Supporting information

Supplementary Tables, Figures and Analyis

## 6. CRediT author statement

**Isabel Hotz:** Conceptualization, Software, Methodology, Validation, Formal analysis, Data curation, Writing - Original Draft, Writing - Review & Editing, Visualization. **Pascal F. Deschwanden:** Methodology, Validation, Formal analysis, Data curation, Writing - Review & Editing. **Franz Liem:** Software, Formal analysis, Investigation, Data curation, Writing - Review & Editing. **Susan Mérillat:** Conceptualization, Investigation, Resources, Data curation, Writing - Review & Editing, Supervision, Project administration, Funding acquisition. **Spyros Kollias:** Validation, Supervision. **Lutz Jäncke:** Conceptualization, Investigation, Resources, Resources, Writing - Review & Editing, Supervision, Project administration, Funding acquisition.

## Acknowledgements

We acknowledge all participants. We are thankful to Susanne Wehrli for her help segmenting WMH masks used in our study, and Christoph Eberle for his valuable help with the graphics.

The current analysis incorporates data from the Longitudinal Healthy Aging Brain (LHAB) database project carried out at the University of Zurich (UZH). The following researchers at the UZH were involved in the design, set-up, maintenance and support of the LHAB database: Anne Eschen, Lutz Jäncke, Franz Liem, Mike Martin, Susan Mérillat, Christina Röcke, and Jacqueline Zöllig.

## Ethics approval

The study was approved by the ethical committee of the canton of Zurich.

## Patient consent statement

Participation was voluntary and all participants gave written informed consent in accordance with the declaration of Helsinki.

## Data and code availability statement

The data for this manuscript are not publicly available because the used consent does not allow for the public sharing of the data. Our code for preparing the data for BIANCA and executing BIANCA can be found here: https://github.com/dynage/bianca

## The following openly available software were used

FreeSurfer (https://surfer.nmr.mgh.harvard.edu/fswiki/DownloadAndInstall)

UBO Detector (https://cheba.unsw.edu.au/research-groups/neuroimaging/pipeline)

BIANCA (https://fsl.fmrib.ox.ac.uk/fsl/fslwiki/BIANCA/Userguide), part of FSL software (RRID: SCR_002823, https://fsl.fmrib.ox.ac.uk) and the MATLAB implementation of LOCATE (https://git.fmrib.ox.ac.uk/vaanathi/LOCATE-BIANCA)

