## Supplementary Tables, Figures and Analyis for "Performance of three freely available methods for extracting white matter hyperintensities: FreeSurfer, UBO Detector and BIANCA"

### Supplementary Table 1

*Table of the articles which validated the methods in this study, listed according to Dice Similarity Coefficient (DSC), sensitivity, and false positive ratio (FPR). This table does not claim to be complete.*

| **Article** | **Algorithm(s)** | **Sample** | **Subjects** | **WMH load of the subjects** | **GS** | **Inter-rater agreement for the GS** | **Type of algorithm validation** | **DSC between GS and algorithm** | **Sensitivity between GS and algorithm** | **FPR between GS and algorithm** | **Additional results between GS and algorithm** | **Population based study** |
|---|---|---|---|---|---|---|---|---|---|---|---|---|
| Ajilore et al. (2014) | **FreeSurfer** | *N = 126*<br>(*n* = 53<br>*n* = 73) | LLD†<br>HC† | ‡ | manually‡<br>(*N* = 20, LLD) | – | cross-sectionally | – | – | – | $r = 0.91$, $p < 10^{-7}$ | – |
| Olsson et al. (2013) | **FreeSurfer** | *N* = 152 | MCI† (incl. dementia) | ‡ | not fully manually – with MRIcron (*N* = 27) | yes (*N* =2)§ | cross-sectionally | – | – | | 2D FLAIR (GS) vs T1w (FS)<br>Spearman's Rho = 0.65<br>ICC = 0.51<br>Kendall's tau = 0.48 § | Gothenburg MCI study |
| Samaille et al. (2012) | **FreeSurfer** | 1) *N* = 24<br>2) *N* = 43 | 1) MCI†,<br>2) CADASIL† | 1) ‡<br>2) very high | 1) manually‡ with anatomist visualization module (brainvisa)<br>2) FLAIR images as base using BioClinica SAS | – | cross-sectionally | 0.40 | – | – | ICC = 0.52 | – |
| Smith et al. (2011) | **FreeSurfer** | *N* = 147<br>(*n* = 40,<br>*n* = 96,<br>*n* = 11) | normal cognition<br>MCI†<br>AD† | ‡ | manually‡<br>(*N* = 10) | _ | cross-sectionally | – | – | – | Intraclass Correlation Coefficient = 0.91 | Community based study |
| Jiang et al. (2018) (developers) | **UBO Detector** | 1) *N* = 400<br>2) *N* = 539 at baseline | 1) + 2) incl. stroke, TIA†, AF†, depression, dementia (not at baseline) | low – high‡ | fully manually (*N* = 40) | – | 1) cross-sectionally<br>2) longitudinally | 0.848 (overall) | 0.913 (overall) | 0.026 (overall) | overall Specificity = 0.989<br>overall Accuracy = 0.989<br>overall OER = 0.224<br>overall DER = 0.039 § | 1) Older Australian Twins Study (OATS)<br>2) Sydney Memory and Ageing Study (Sydney MAS) |

| | | | | | | | | | | | | |
|---|---|---|---|---|---|---|---|---|---|---|---|---|
| Griffanti et al. (2016) (developers) | **BIANCA** | 1) *N* = 85<br>2) *N* = 474 | 1) AD†, MCI†, subjective CI†, HC†<br>2) (neurodegenerative cohort = NDGEN) non-disabling stroke, TIA (vascular cohort = OXVASC) | medium‡ | fully manually<br><br>1) *N* = 21<br>2) *N* = 109 | _ | cross-sectionally | 1) 0.76<br>2) 0.52 | _ | 1) 0.22<br>2) 0.46 | 1) ICC = 0.990, DER[a] = 0.03, OER[a] = 0.46¶<br><br>2) ICC = 0.919, DER[a] = 0.19, OER[a] = 0.76¶ | 1) Oxford Project to Investigate Memory and Ageing (OPTIMA)<br>2) Oxford Vascular Study (OXVASC) |
| Ling et al.(2018) | **BIANCA** | 1) *N* = 90 (2D FLAIR<br>2) *N* = 66/90 (3D FLAIR) | CADASIL† | very high | semi-automated (*N* = 20) | yes (*N* =2)§ | cross-sectionally | 1) median 0.79<br>2) median 0.76 | _ | 1) 0.23<br>2) 0.20 | 1) ICC = 0.81¶<br>2) ICC = 0.78¶ | CADASIL study |
| Sundaresan et al. (2018) (developers) | **LOCATE** | 1) *N* = 21<br>2) *N* = 18<br>3) *N* = 15<br>4) *N* = 19<br>5) *N* = 60 | 1) + 2) same as in Griffanti et al. (2018)<br>3) CADASIL†<br>4) HC<br>5) from the WMH segmentation study MWSC, see Kuijf et al. (2019) | medium‡<br>–<br>very high | 1) + 2) see Griffanti et al. (2018)<br>3) no GS<br>4) no GS<br>5) fully manually (Kuijf et al., 2019)<br><br>For 3) + 4) they used BIANCA trained with 2) as a reference – no manually traced GSs | 5) yes (N =2)§ | cross-sectionally | 1) 0.77<br>2) 0.75<br>3) 0.79<br>4) –<br>5) range = 0.63 – 0.73 | 1) 0.03 (increase to global threshold of BIANCA)<br>2) 0.10 (increase)<br>3) 0.48 (increase)<br>4) –<br><br>1), 2), 5) for PVWMH and DWMH‡ | 1) 0.001(increase)<br>2) 0.002 (increase)<br>3) 0.00<br>4) –<br>5) small increase§ | – | – |
| (Vanderbecq et al., 2020) | **BIANCA & UBO Detector (and 5 others)** | 1) *N* = 147<br>2) *N* = 60 | 1) AD, MCI, CN† (Research dataset)<br>2) Cognitive impairment (Clinical routine dataset, incl. 10 images with artifacts) | medium – high | not fully manually – with ITK-SNAP<br><br>1) *N* = 20<br>2) *N* = 20 | yes (*N* =2)§ | cross-sectionally | BIANCA<br>1) 0.469<br>2) 0.607<br><br>UBO<br>1) 0.486<br>2) 0.560 | – | BIANCA<br>1) 0.393<br>2) 0.404<br><br>UBO<br>1) 0.587<br>2) 0.471 | BIANCA<br>1) ICC = 0.417<br>2) ICC = 0.859<br>1) FNR = 0.481<br>2) FNR = 0.296¶<br><br>UBO<br>1) ICC = 0.881<br>2) ICC = 0.734<br>1) FNR = 0.360<br>2) FNR = 0.353¶ | Alzheimer's Disease Neuroimaging Initiative (ADNI) database |

*Notes:* GS = gold standard; Tp = Time point; *N* = Number of subjects; ICC = Interclass Correlation Coefficient, OER = Outline Error Rate.
† Subjects' clinical status: MDD = major depression disorder, LLD = late-life depression, HC = healthy controls, MCI = amnestic mild cognitive impairment; CADASIL = Cerebral Autosomal Dominant Arteriopathy with Subcortical Infarcts and Leukoencephalopathy, AD = Alzheimer's disease, TIA = transient ischaemic attack, AF = arterial fibrillation, CI = cognitive impairment.
‡no clear information; §The number of *N* takes into account the number of operators included in the calculation for the inter-operator reliability, ¶ for more results/information see article; ¶
If multiple results are given, those based on the highest DSC are given.

### Supplementary Table 2

***Validation of the gold standards** for 3D FLAIR (n = 16) and 2D FLAIR (n = 10) images including: Dice Similarity Coefficient (DSC), Detection Error Rate (DER), Hausdorff Distance for the 95th percentile (H95), Outline Error Rate (OER), Interclass Correlation Coefficient (ICC) according to number (N) of subjects with low (< 5cm$^3$), medium (5 – 15 cm$^3$), and high (> 15 cm$^3$) mean WMH load for manual segmentation, and mean inter-rater agreements. Mean WMH volumes for the T1w (n = 16) gold standard images according to WMH load.*

| | | | | **Overlap agreement** | | **Resemblance agreement** | **Volumetric agreement** | |
|---|---|---|---|---|---|---|---|---|
| **Image** | ***N*** | **Volume in cm$^3$ ± SD†** | **Load** | **DSC** | **DER** | **H95[b]** | **OER** | **ICC** |
| 3D FLAIR | 7 | 3.88 ± 0.62 | low | 0.70 | | | | |
| 3D FLAIR | 7 | 9.28 ± 3.08 | medium | 0.73 | | | | |
| 3D FLAIR | 2 | 20.44 ± 2.95 | high | 0.87 | | | | |
| **Total** | **16** | **8.31 ± 5.81** | **medium** | **0.73 ± SD 0.09** | **0.12 ± SD 0.08** | **4.21 ± SD 3.20** | **0.30 ± SD 0.13** | **0.96***** |
| 2D FLAIR | 3 | 3.32 ± 1.07 | low | 0.63 | | | | |
| 2D FLAIR | 7 | 8.11 ± 3.22 | medium | 0.73 | | | | |
| **Total** | **10** | **6.68 ± 3.54** | **medium** | **0.67 ± SD 0.11** | **0.15 ± SD 0.12** | **7.19 ± SD 6.14** | **0.51 ± SD 0.16** | **0.82**** |
| T1w | 7 | 4.25 ± 0.39 | low | - | | | | |
| T1w | 7 | 7.86 ± 2.21 | medium | - | | | | |
| T1w | 2 | 19.85 ± 0.49 | high | - | | | | |
| **Total** | **16** | **7.78 ± 5.22** | **medium** | - | - | - | - | - |

*Notes:* †Mean WMH volume of the gold standards in cm$^3$ ± standard deviation (SD). [b] H95 in mm. In the T1w sequences the images were segmented by one operator.
*** $p < 0.001$; ** $p = 0.0012$

### Supplementary Table 3

***Validation of the Fazekas scale.** Mean inter-operator reliabilities using a weighted Cohen's kappa (Cohen, 1968) between three operators over four time points (Tp1, Tp2, Tp3, Tp5)(N = 800; p < 0.001).*

| **Operator** | **Total WMH** | **PVWMH** | **DWMH** |
|---|---|---|---|
| **O1 vs O2** | κ = 0.769 | κ = 0.654 | κ = 0.764 |
| **O2 vs O3** | κ = 0.887 | κ = 0.858 | κ = 0.811 |
| **O3 vs O1** | κ = 0.809 | κ = 0.700 | κ = 0.812 |

O = Operator; κ = Cohen's Kappa; Total WMH = Total White Matter Hyperintensities; PVWMH = Periventricular White Matter Hyperintensities; DWMH = Deep White Matter Hyperintensities.
Cohen's kappa: no agreement: ≤ 0, none to slight agreement: 0.01 – 0.20, fair agreement: 0.21 – 0.40, moderate agreement: 0.41 – 0.60, substantial agreement: 0.61 – 0.80, almost perfect agreement: 0.81 – 1.00 (Landis & Koch, 1977).

### Supplementary Table 4

***Description of the Fazekas scale rating of all three operators.*** *Displayed are the number and percentage of distributed Fazekas scores divided into DWMH, PVWMH and total WMH per time point (Tp).*

| **Time point (TP)** | **DWMH** | | | **PVWMH** | | | **Total WMH** | | |
|---|---|---|---|---|---|---|---|---|---|
| | **Fazekas score** | ***n*** | **%** | **Fazekas score** | ***n*** | **%** | **Fazekas score** | ***n*** | **%** |
| **Tp 1 (*N* = 231)** | 0 | 25 | 10.823 | 0 | 11 | 4.762 | 0 | 5 | 2.165 |
| | 1 | 154 | 66.667 | 1 | 91 | 39.394 | 1 | 20 | 8.658 |
| | 2 | 45 | 19.481 | 2 | 112 | 48.485 | 2 | 75 | 32.468 |
| | 3 | 7 | 3.030 | 3 | 17 | 7.359 | 3 | 77 | 33.333 |
| | | | | | | | 4 | 40 | 17.316 |
| | | | | | | | 5 | 9 | 3.896 |
| | | | | | | | 6 | 6 | 2.597 |
| **Tp 2 (*N* = 207)** | 0 | 16 | 7.729 | 0 | 8 | 3.865 | 0 | 2 | 0.966 |
| | 1 | 140 | 67.633 | 1 | 78 | 37.681 | 1 | 16 | 7.729 |
| | 2 | 43 | 20.773 | 2 | 105 | 50.725 | 2 | 63 | 30.435 |
| | 3 | 8 | 3.865 | 3 | 16 | 7.729 | 3 | 77 | 37.198 |
| | | | | | | | 4 | 32 | 15.459 |
| | | | | | | | 5 | 12 | 5.797 |
| | | | | | | | 6 | 5 | 2.415 |
| **Tp 3 (*N* = 196)** | 0 | 12 | 6.122 | 0 | 7 | 3.571 | 0 | 3 | 1.531 |
| | 1 | 135 | 68.878 | 1 | 71 | 36.224 | 1 | 11 | 5.612 |
| | 2 | 41 | 20.918 | 2 | 100 | 51.020 | 2 | 58 | 29.592 |
| | 3 | 8 | 4.081 | 3 | 18 | 9.184 | 3 | 77 | 39.286 |
| | | | | | | | 4 | 28 | 14.286 |
| | | | | | | | 5 | 14 | 7.143 |
| | | | | | | | 6 | 5 | 2.551 |
| **Tp 5 (*N* = 166)** | 0 | 9 | 5.422 | 0 | 7 | 4.217 | 0 | 2 | 1.205 |
| | 1 | 117 | 70.482 | 1 | 57 | 34.337 | 1 | 9 | 5.422 |
| | 2 | 32 | 19.277 | 2 | 87 | 52.410 | 2 | 49 | 29.518 |
| | 3 | 8 | 4.819 | 3 | 15 | 9.036 | 3 | 67 | 40.361 |
| | | | | | | | 4 | 22 | 13.253 |
| | | | | | | | 5 | 12 | 7.229 |
| | | | | | | | 6 | 5 | 3.012 |
| **ALL Tp's (*N* = 800)** | 0 | 62 | 7.750 | 0 | 33 | 4.125 | 0 | 12 | 1.500 |
| | **1** | **546** | **68.250** | 1 | 297 | 37.125 | 1 | 56 | 7.000 |
| | 2 | 161 | 20.125 | **2** | **404** | **50.500** | **2** | **245** | **30.625** |
| | 3 | 31 | 3.875 | 3 | 66 | 8.250 | **3** | **298** | **37.250** |
| | | | | | | | **4** | **122** | **15.250** |
| | | | | | | | 5 | 47 | 5.875 |
| | | | | | | | 6 | 20 | 2.500 |

Tp = Time point; DWMH = deep white matter hyperintensities; PVWMH = periventricular white matter hyperintensities; total WMH = total white matter hyperintensities.

### Supplementary Table 5

***Threshold and k optimization in UBO Detector** with the sixteen 3D and 2D FLAIR gold standards (+ T1w) using the leave-one out cross-validation method for k 3 and 5, and the thresholds 0.7 and 0.9.*

| Modality | Number of NN | Threshold = 0.7 | | Threshold = 0.9 | |
|---|---|---|---|---|---|
| 3D FLAIR | ***k* = 3** | DSC = | 0.4793 | DSC = | 0.4793 |
| | | OER = | 0.6943 | OER = | 0.6943 |
| | | DER = | 0.3471 | DER = | 0.3471 |
| | | H95 = | 12.7643 | H95 = | 12.7634 |
| | | ICC = | 0.852 | ICC = | 0.852 |
| | ***k* = 5** | DSC = | 0.4998 | DSC = | 0.4521 |
| | | OER = | 0.6872 | OER = | 0.6421 |
| | | DER = | 0.3131 | DER = | 0.4536 |
| | | H95 = | 11.5331 | H95 = | 16.0401 |
| | | ICC = | 0.876 | ICC = | 0.826 |
| 2D FLAIR | ***k* = 3** | DSC = | 0.5307 | DSC = | 0.5310 |
| | | OER = | 0.6112 | OER = | 0.6113 |
| | | DER = | 0.3275 | DER = | 0.3267 |
| | | H95 = | 13.5472 | H95 = | 13.5216 |
| | | ICC = | 0.928 | ICC = | 0.927 |
| | ***k* = 5** | DSC = | 0.5283 | DSC = | 0.5281 |
| | | OER = | 0.6105 | OER = | 0.5976 |
| | | DER = | 0.3329 | DER = | 0.3461 |
| | | H95 = | 15.4510 | H95 = | 14.1409 |
| | | ICC = | 0.919 | ICC = | 0.926 |

*NN* = nearest neighbor; DSC = mean DSC; OER = Outline Error Rate; DER = Detection Error Rate; H95 = Hausdorff Distance for the 95 percentile; ICC = Intraclass-Correlation Coefficient.

### Supplementary Table 6

***Threshold optimization in BIANCA** with the 16 3D and 2D FLAIR gold standards (+ T1w) using the leave-one out cross-validation method for the thresholds 0.90, 0.95, and 0.99.*

| modality | Threshold = 0.90 | | Threshold = 0.95 | | Threshold = 0.99 | |
|---|---|---|---|---|---|---|
| 3D FLAIR | DSC = | 0.5186 | DSC = | 0.5738 | DSC = | **0.6015** |
| | OER = | 0.7729 | OER = | 0.6713 | OER = | 0.6353 |
| | DER = | 0.1899 | DER = | 0.1811 | DER = | 0.1617 |
| | H95 = | 9.9126 | H95 = | 8.1674 | H95 = | 6.1988 |
| | ICC = | 0.169 | ICC = | 0.332 | ICC = | 0.743 |
| 2D FLAIR | DSC = | 0.4519 | DSC = | 0.5190 | DSC = | **0.5607** |
| | OER = | 0.8904 | OER = | 0.7488 | OER = | 0.6636 |
| | DER = | 0.2058 | DER = | 0.2132 | DER = | 0.2149 |
| | H95 = | 10.0643 | H95 = | 9.0159 | H95 = | 8.4429 |
| | ICC = | 0.332 | ICC = | 0.551 | ICC = | 0.859 |

DSC = mean DSC; OER = Outline Error Rate; DER = Detection Error Rate;
H95 = Hausdorff Distance for the 95 percentile; ICC = Intraclass-Correlation Coefficient

### Supplementary Table 7

***Additional accuracy metrics calculated using BIANCA's evaluations script*** *to better compare our data to those from BIANCA's original study.*

| Modality | DSC | FDR | FNR | FDR (Cluster) | FNR (Cluster) |
|---|---|---|---|---|---|
| **2D FLAIR** | 0.561 | 0.369 | 0.425 | 0.662 | 0.470 |
| **3D FLAIR** | 0.602 | 0.333 | 0.390 | 0.701 | 0.371 |

DSC = Dice Similarity Coefficient; FDR = false discovery rate; FNR = false negative rate; FDR (Cluster) = cluster-level false discovery rate (sensitivity of individual lesions); FNR (Cluster) = cluster-level false negative rate.

### Supplementary Table 8

***Mean WMH volume differences relative to the respective gold standard*** *in $cm^3$ and* ***mean absolute WMH volume differences*** *in $cm^3$ relative to the respective gold standard for the different algorithms and modalities (n = 16).*

| | Mean volume difference to gold standard | Rank | Mean absolute volume difference to gold standard | Rank |
|---|---|---|---|---|
| **FreeSurfer (T1w)** | 4.345 | 5 | 4.345 | 5 |
| **UBO (2D FLAIR)** | 0.478 | 3 | 2.000 | **1** |
| **UBO (3D FLAIR)** | 2.310 | 4 | 2.356 | **2** |
| **BIANCA (2D FLAIR)** | 0.337 | **2** | 3.197 | 4 |
| **BIANCA (3D FLAIR)** | 0.005 | **1** | 2.612 | 3 |
| **ANOVA** | F = 5.927, $p < 0.001$ | | F = 2.105, $p < 0.089$ | |
| **Post-Hoc (Tukey)** | FreeSurfer > BIANCA (2D FLAIR), BIANCA (3D FLAIR), UBO (2D FLAIR) | | No differences | |

*Notes:* Input modality for the algorithms: FreeSurfer: T1w, UBO Detector and BIANCA: 3D FLAIR + T1w and 2D FLAIR + T1w. The rank of FreeSurfer does not change when considering the absolute mean differences to the gold standard. BIANCA 3D and 2D FLAIR must however give up his 1 and 2 rank respectively to UBO 2D and 3D FLAIR.

**Supplementary Table 9** demonstrates, that the DSC – calculated from all algorithm's outputs – is sensitive to the WMH load, categorized according to the respective manually segmented gold standard per modality (T1w, 3D FLAIR, 2D FLAIR). However, because of the constant underestimation of the mean WMH volumes by FreeSurfer, it already reached the maximum DSC at the medium WMH load. Therefore, FreeSurfer could not achieve an improvement of the DSC from medium to high WMH exposure in terms of DSC.

**Supplementary Table 9**

*Comparison of the mean Dice Similarity Coefficient (DSC) of the three algorithms (FreeSurfer T1w, UBO Detector and BIANCA 3D FLAIR + T1w and 2D FLAIR + T1w) according to subjects with low (L) (< 5cm$^3$), medium (M) (5 – 15 cm$^3$), and high (H) (> 15 cm$^3$) mean WMH load for the corresponding manual segmentation of the same 16 subjects per modality (T1w, 3D FLAIR, 2D FLAIR).*

| **WMH load GS** | | **T1-w †** | **3D FLAIR ‡** | | **2D FLAIR §** | |
|---|---|---|---|---|---|---|
| | | **FreeSurfer** | **UBO Detector** | **BIANCA** | **UBO Detector** | **BIANCA** |
| | | **Mean DSC** | | | | |
| **< 5 cm$^3$** | **L** | 0.371 | 0.414 | 0.501 | 0.432 | 0.482 |
| **5 – 15 cm$^3$** | **M** | 0.487 | 0.550 | 0.674 | 0.556 | 0.577 |
| **> 15 cm$^3$** | **H** | 0.473 | 0.638 | 0.700 | 0.668 | 0.684 |
| **Total ± SD** | | 0.434 ± 0.111 | 0.501 ± 0.124 | 0.602 ± 0.133 | 0.531 ± 0.113 | 0.561 ± 0.118 |

*Notes:* † T1-w gold standard: L: (*n* = 7), M: (n = 7), H: (*n* = 2). ‡ 3D FLAIR gold standard: L: (*n* = 7), M: (*n* = 7), H: (*n* = 2).

### Detailed description of post-hoc outlier analysis

**In a first step,** we determined how many comparisons between two measurement points were possible for each subject in the entire data set. We differentiated between 1-year intervals (for example: baseline – 1-year follow-up / 1-year follow-up – 2-year follow-up), 2-year-intervals, 3-year-intervals, and 4-year-intervals. For BIANCA 2D 209 subjects provide data over time (at least two time points). These 2D FLAIR images (*N* = 762 images) allowed 531 intervals between two measurement points (1-year intervals: *n* = 369, 2-year intervals: *n* = 145 etc.). Importantly, overlapping intervals were not included. If a subject had data for the baseline, 1-year and 2-year follow-up, only the comparisons «baseline vs. 1-year follow-up» and «1-year follow-up vs. 2-year follow-up» were considered. The comparison «baseline vs. 2-year follow-up» was only considered if a subject missed 1-year follow-up data.

**Secondly,** we calculated the percent WMH volume change from the previous to the subsequent measurement point for each subject, for each combination of algorithm and modality (e.g., BIANCA 2D), and for the available time interval (1-year interval). Then, for each algorithm-by-modality combination and available time interval. The mean percent change was determined as well as the standard deviation (SD). These metrics were averaged across the five algorithm-by-modality combinations and used to calculate a general «range of tolerance», in which WMH volume increase and decrease for a given time interval were considered representing true change. The upper limit of the increases (average across algorithm-by-modality + 1SD) and the lower limit of the decreases (average across algorithm-by-modality - 1SD) were used to identify the outlier. **Supplementary Table 10** lists the algorithm- and modality-specific WMH volume changes in percent, the mean WMH volume changes across all algorithms, and the associated «range of tolerance» for all time intervals. WMH volume increases and decreases outside of these «ranges of tolerance» were considered as outlier. **Table 7** provides information on the number of outlier changes per algorithm and modality in comparison to the number of possible comparisons between two measurement points. This table also contains information on the distribution of outlier changes in dependence of WMH load (low, medium, high) – using the Fazekas scale – and informs about the percentage of subjects, in which outlier WMH volume changes occurred. The categories were divided into the following categories: Fazekas score 0 – 2 = low WMH load, Fazekas score 3 and 4 = medium WMH load, and Fazekas score 5 and 6 = high WMH load. In each case, the previous measurement point, in the individual change trajectories over time, served as the basis for the classification.

### Supplementary Table 10

*Algorithm- and modality-specific WMH volume changes in percent (± 1SD), mean WMH volume changes across all algorithms (± 1SD), and the associated «range of tolerance» per interval (1-, 2-, 3-, 4- years) in the output of dataset 1 (FreeSurfer = FS), dataset 2 (UBO 2D, BIANCA 2D) and dataset 3 (UBO 3D, BIANCA 3D). The «range of tolerance» (in bold) was defined by the mean plus/minus the standard deviation (± 1 SD) of increase and decrease.*

| Interval | FS | UBO 3D | UBO 2D | BIANCA 3D | BIANCA 2D | Mean and ± § 1 SD in % | «Range of tolerance»[c] |
|---|---|---|---|---|---|---|---|
| | Mean in % (± 1 SD in %) | | | | | | |
| **1 Year** | 6.24% (±10.26%) *N* = 397 | 6.59% (±12.99%) *N* = 12 | 8.64% (±19.42%) *N* = 360 | 49.96% (±157.30%) *N* = 12 | 31.85% (±76.45%) *N* = 369 | 20.66% (±55.22%) | |
| **Increase †** | 9.87% (+9.15%) *N* = 297 | 14.31% (+4.82%) *N* = 8 | 17.54% (+16.95%) *N* = 241 | 76.82% (+174.65%) *N* = 9 | 64.80% (+76.92%) *N* = 236 | 36.67% (+56.50%) | -19.83%; **93.17% –** |
| **Decrease ‡** | -4.54% (-3.75%) *N* = 100 | -8.84% (-9.37%) *N* = 4 | -9.45% (-8.47%) *N* = 119 | -30.63% (-33.06%) *N* = 3 | -26.61% (-19.41%) *N* = 133 | -16.01% (-14.81%) | **-30.82%**; 1.2% |
| **2 Years** | 12.53% (±14.83%) *N* = 170 | 19.06% (±20.34%) *N* = 21 | 16.46% (±21.98%) *N* = 145 | 41.04% (±82.43) *N* = 21 | 29.34% (±74.98%) *N* = 145 | 23.68% (±42.91%) | |
| **Increase †** | 16.86% (+12.89%) *N* = 138 | 24.33% (+15.86%) *N* = 18 | 23.05% (+18.91%) *N* = 117 | 50.35% (+85.7%) *N* = 18 | 66.91% (+73.27%) *N* = 88 | 36.30% (+41.33%) | -5.03%; **77.63% –** |
| **Decrease ‡** | -6.13% (-4.77%) *N* = 32 | -12.59% (-15.64%) *N* = 3 | -11.06% (-8.11%) *N* = 28 | -14.84% (-7.13%) *N* = 3 | -28.68% (-21.68%) *N* = 57 | -14.66% (-11.47%) | **-26.13%**; 3.19% |
| **3 Years** | 22.76% (± 8.10%) *N* = 2 | 27.96% (± 34.14%) *N* = 15 | 16.12% (± 20.73%) *N* = 16 | 40.79% (± 43.09%) *N* = 15 | 55.95% (± 73.40%) *N* = 16 | 32.72% (±3 5.89%) | |
| **Increase †** | 22.76% (+8.10%) *N* = 2 | 33.81% (+32.88%) *N* = 13 | 21.59% (+18.95%) *N* = 13 | 48.14% (+41.53%) *N* = 13 | 92.12% (+58.21%) *N* = 11 | 43.68% (+31.93%) | 11.75%; **75.61% –** |
| **Decrease ‡** | *N* = 0 | -10.07% (-2.88%) *N* = 2 | -7.57% (-5.55%) *N* = 3 | -6.98% (-5.27%) *N* = 2 | -23.62% (-14.82%) *N* = 5 | -12.06% (-7.13%) | **-19.19**; 4.92% |
| **4 Years** | – | – | *N* = 1 | – | *N* = 1 | ¶ | ¶ |

*Notes:* † Increase of WMH volume in a specific annual interval per algorithm output; ‡ Decrease of WMH volume in a specific annual interval per algorithm output; § The upper limit of the increases (equally weighted mean + 1SD) and the lower limit of the decreases (equally weighted mean - 1SD) were used to identify the suspicious cases.
¶ Due to missing data no «Range of tolerance» could be calculated for the 4-year interval.

### Supplementary Table 11

*Display per subsets and per algorithm (BIANCA 2D, BIANCA 3D, UBO 2D, UBO 3D, FreeSurfer T1w) according the WMH load of the total number (N) with the outputted number (N) of segmented scans, number (n) of subjects with longitudinal data (at least 2 time points), number and percentage (in brackets) of subjects with outlier in longitudinal data, number of intervals between two measurement points, and number and percentage (in brackets) of outlier intervals between two measurement points.*

| Subsets / WMH load | N of segmented scans | n of subjects | n (and %) of subjects with outlier in longitudinal data | | n of intervals † between two measurement points | n (and %) of outlier intervals between two measurement points | |
|---|---|---|---|---|---|---|---|
| **BIANCA 2D** | 762 | 209 | **109** | **(52.15%)** | **531** | **161** | **(30.32%)** |
| low | | | | | | 90 | (55.90%) |
| medium | | | | | | 66 | (40.99%) |
| high | | | | | | 5 | (3.11%) |
| **BIANCA 3D** | 166 | 39 | **7** | **(17.95%)** | **48** | **8** | **(16.67%)** |
| low | | | | | | 3 | (37.50%) |
| medium | | | | | | 5 | (62.50%) |
| high | | | | | | 0 | (0%) |
| **UBO 2D** | 757 ‡ | 209 | **7** | **(3.35%)** | **523** | **7** | **(1.34%)** |
| low | | | | | | 4 | (57.14%) |
| medium | | | | | | 1 | (14.29%) |
| high | | | | | | 2 | (28.57%) |
| **UBO 3D** | 166 | 39 | **2** | **(5.13%)** | **48** | **2** | **(4.17%)** |
| low | | | | | | 0 | (0%) |
| medium | | | | | | 2 | (100.00%) |
| high | | | | | | 0 | (0%) |
| **FreeSurfer T1w** | 800 | 213 | **0** | **(0%)** | **569** | **0** | **(0%)** |
| low | | | | | | 0 | (0%) |
| medium | | | | | | 0 | (0%) |
| high | | | | | | 0 | (0%) |

*Notes*: WMH load is divided into low, medium, and high WMH load according to the Fazekas scale. Fazekas score 0 – 2 = low WMH load, Fazekas score 3 and 4 = medium WMH load, and Fazekas score 5 and 6 = high WMH load.
† Explanation «intervals between two measurement points»: If a subject had 3 time points (Tp1, Tp2 and Tp5) this would result in two existing intervals. ‡ The data point with the segmentation error (segmented eyeballs) is included.

### Supplementary Figures

#### Supplementary Figure 1

***Data structure and study design*** *divided by number of scans, time points (Tps), and modality (T1w, 2D FLAIR, 3D FLAIR). The* ***Fazekas scale*** *was applied to validate FreeSurfer, UBO Detector, and BIANCA algorithms using longitudinal data. The longitudinal data were also used to associate chronological age with the algorithms outputs and to do a post-hoc outlier analysis. The* ***manual gold standards*** *were used for cross-sectional comparisons of segmentations accuracy, WMH volumes, and as a training dataset for BIANCA.*

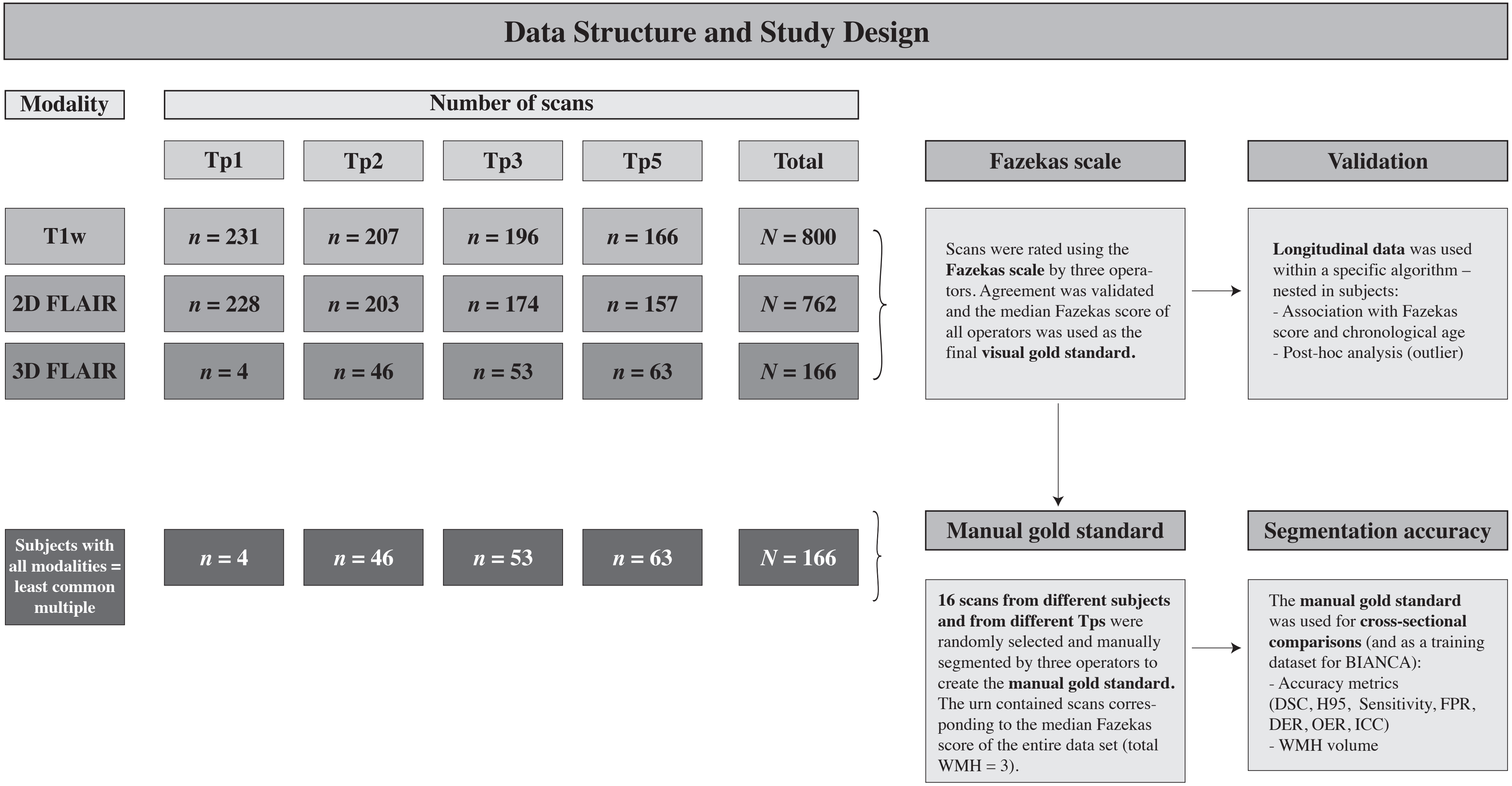

### Supplementary Analysis

### Comparison of the threshold methods: BIANCA and LOCATE

LOCally Adaptive Threshold Estimation (LOCATE) is a method that, in contrast to a global threshold, determines spatially adaptive thresholds in different regions in the probability map. This allows to overcome the influence of spatial heterogeneity of lesion probabilities due to changes in lesion contrast, load and distribution on the final threshold map of WMH. As input, LOCATE uses the lesion probability map at subject level obtained from a WMH detection algorithm. For more details on the descriptions see Sundaresan et al. (2018). Before applying LOCATE, we normalized the FLAIR images' values within the brain masks to a range of *0* to *1* to avoid different ranges of intensities between the images.

##### Comparison of the threshold methods

In order to determine whether BIANCA with the best global threshold of 0.99 or the LOCAL method is more suitable for our data, for each method we carried out a leave-one-out cross-validation against both the 3D FLAIR and 2D FLAIR gold standards.

As shown in **Supplementary Table 12** on average BIANCA marks significantly less false positives compared to LOCATE. In BIANCA with the 3D FLAIR + T1w images input, the mean H95 was significantly better than in LOCATE with the same input. Only the mean sensitivity was better in LOCATE than in BIANCA. The mean WMH volume of LOCATE was significantly different from the gold standard and also significantly higher than the mean WMH volume of BIANCA. This was also reflected in the low and non-significant ICC(3,1) of LOCATE.

Based on the results in **Supplementary Table 12**, the outliers of each DSC in LOCATE compared to the gold standard (DSC range: LOCATE 2D FLAIR + T1w = 0.194 – 0.687, 3D FLAIR + T1w = 0.165 – 0.705; BIANCA 2D FLAIR + T1w = 0.334 – 0.734, BIANCA 3D + T1w = 0.292 – 0.783), and the visual inspections, we decided to use BIANCA for the entire sample.

### Supplementary Table 12

***Statistical comparison between BIANCA (global threshold of 0.99) and LOCATE*** *– using the gold standard with the sixteen subjects per modality (3D FLAIR + T1w, 2D FLAIR + T1w) as reference – with the different accuracy metrics, the WMH volumes. Further a comparison of the WMH volumes and the ICCs from the automated methods and the manually segmented gold standards.*

| | **BIANCA 3D FLAIR†** | **LOCATE 3D FLAIR[b]** | **WILCOXON** | | **BIANCA 2D FLAIR[a]** | **LOCATE 2D FLAIR[b]** | **WILCOXON** | |
|---|---|---|---|---|---|---|---|---|
| | | | ***W*** | ***p*** | | | ***W*** | ***p*** |
| **DSC** | 0.602 | 0.552 | 144 | 0.564 | 0.561 | 0.512 | 150 | 0.423 |
| **OER** | 0.635 | 0.706 | 124 | 0.897 | 0.664 | 0.751 | 106 | 0.423 |
| **DER** | 0.162 | 0.190 | 87 | 0.128 | 0.215 | 0.224 | 117 | 0.696 |
| **H95** | 6.200 | 9.298 | 54 | 0.004 | 8.443 | 9.314 | 102 | 0.337 |
| **FP** | 5619.6 | 19241.4 | 46 | 0.001 | 3512.6 | 10820.1 | 46 | 0.001 |
| **TP** | 9939.1 | 12761.4 | 99 | 0.287 | 5937.1 | 7605.6 | 97 | 0.254 |
| **FPR** | **0.0003** | 0.0009 | 46 | 0.001 | **0.0004** | 0.0011 | 46 | 0.001 |
| **Sensitivity** | 0.611 | **0.812** | 19 | < 0.001 | 0.575 | **0.766** | 32 | < 0.001 |
| | **COEFF *p*-value** | **COEFF *p*-value** | | | **COEFF *p*-value** | **COEFF *p*-value** | | |
| **Vol. method vs Vol. GS[a]** | *W* = 133 *p* = 0.867 | W = 198 *p* = 0.007 | | | *W* = 123 *p* = 0.867 | W = 194 *p* = 0.012 | | |
| **ICC(3,1)** | 0.743 < 0.001 | 0.206 *p* = 0.141 | | | 0.859 < 0.001 | 0.526 *p* < 0.05* | | |
| **Vol. in cm³** | 8.317 | 17.106 | | | 8.695 | 16.955 | | |
| **Vol. BIANCA vs Vol. LOCATE[a]** | ***W* = 65 *p* = 0.017** | | | | ***W* = 64 *p* = 0.015** | | | |

*Notes:* †Comparison by Wilcoxon-rank-sum-test.
DSC = mean DSC; OER = Outline Error Rate; DER = Detection Error Rate; H95 = Hausdorff distance for the 95 percentile; FP = false positives; TP = true positives; FPR = false positive rate; Vol. = WMH Volume; ICC = Interclass Correlation Coefficient
